# Multiple Fault Analysis and Drug Therapy on Signaling Pathways Using Dynamic Bayesian Network-based Model

**DOI:** 10.64898/2026.06.11.731601

**Authors:** Tapan Chowdhury, Ekarsi Lodh, Shalini Majumder, Anindya Maitra, Anjali Agarwal, Arundhuti Sur, Sohini Sarkar

## Abstract

Cancer-associated signaling pathways often exhibit abnormal activation under simultaneous dysregulation of multiple molecular components. This study presents a probabilistic temporal Dynamic Bayesian Network (DBN)-based framework for analyzing multi-fault behaviour and intervention response in Growth Factor (GF) and Mitogen-Activated Protein Kinase (MAPK) signaling pathways. Unlike deterministic Boolean propagation, the proposed model represents each pathway component through an activation probability and propagates these probabilities over discrete time steps using soft-logic update rules. One-, two-, three-, and four-fault scenarios were systematically evaluated under a common lowest-burden input vector. The resulting output probabilities were summarized using an encoded pathway-burden score, and known-drug combinations were ranked using efficiency scores relative to no-intervention baselines. Pareto analysis was further used to balance intervention efficiency against drug-vector burden, while a custom dual-target search was performed to identify computational intervention hypotheses beyond predefined drug targets. Results showed that encoded burden increased with fault order in both pathways, with MAPK producing a higher baseline burden than GF. Among known-drug vectors, U0126+LY294002+Temsirolimus consistently emerged as the strongest low-burden candidate, achieving efficiency close to the maximum six-drug vector. Custom dual-target analysis identified ERK1/2+RPS6KB1 in GF and Raf+MEK1 in MAPK as high-impact computational target pairs. Runtime benchmarking showed that batched vectorized NumPy execution substantially improved scalability for higher-order fault simulations. Overall, the framework provides an interpretable and scalable approach for probabilistic pathway-level fault analysis and intervention prioritization.

## 1 Introduction

Cell growth is controlled by a set of complex signaling cascades, which basically involve protein-protein interactions that translate signals from growth factors to transcription factors or adaptors [1]. Abnormality in these signal transduction mechanisms is often associated with uncontrolled cellular proliferation, which ultimately leads to cancer [2]. This study utilizes Dynamic Bayesian Networks (DBNs) [3] to capture and emulate such deviations. This research specifically targets the Growth Factor (GF) and Mitogen-Activated Protein Kinase (MAPK) [4], [5], [6], [7], [8] signaling networks. By assuming the binary nature of proteins, either active or inactive (0), DBNs offer an effective modeling approach. Particularly, the PI3K/AKT/mTOR and Ras/MEK/ERK (MAPK) signaling networks [4] are extensively examined in the context of conditions, such as breast cancer. These networks feature intricate communications among proteins, each with input, function, and output components. The output of one protein-protein interaction serves as the input for another, ultimately determining whether cell division (mitosis) or cell death (apoptosis) is triggered [9]. The modeling procedure aids in simulating circumstances that result in cancer [7]. It has been demonstrated that under specific drug combinations, the EGFR family of protein receptors within the network may reduce the effects of preexisting malfunctions [9], [10].

However, several pathway-level therapeutic modeling studies have relied on Boolean or digital-circuit representations of signaling pathways, often evaluating fault effects through discrete stuck-at perturbations and drug-response logic [7], [9], [11]. While such logic-based models are useful abstractions, binary on/off representations may oversimplify biochemical signaling, where interactions can show graded, probabilistic, and context-dependent behaviour [12], [13], [14]. In this study, we extend DBN-based signaling pathway analysis by representing each node through an activation probability and by evaluating the effect of multiple concurrent faults in the GF and MAPK pathways. The final pathway response is summarized using an encoded burden score, which enables comparison between fault-only and intervention-treated states.

In addition to simulating one, two, three, and four-fault scenarios, this study evaluates all 128 combinations of seven known drugs mapped to pathway-specific intervention points. Candidate interventions are ranked using efficiency scores and Pareto-based analysis so that both burden reduction and intervention complexity can be considered. Furthermore, custom dual-target interventions are explored as computational hypotheses for identifying pathway-node pairs that may reduce downstream output burden beyond the predefined known-drug target set.

The main contributions of this study are as follows:

i. A probabilistic temporal DBN framework is developed for modeling GF and MAPK signaling pathway behaviour under fault-free, multi-fault, and intervention-treated conditions.
ii. Multi-fault dysregulation is systematically evaluated across one-, two-, three-, and four-fault scenarios.
iii. An encoded pathway-burden measure and efficiency-score formulation are used to compare known-drug vectors across fault combinations.
iv. Pareto-based ranking is applied to identify drug vectors that balance intervention efficiency and intervention burden.
v. A custom dual-target search is performed to identify computational inhibition-point hypotheses beyond the predefined known-drug target set.

The remainder of the paper is organized as follows. Section 2 reviews related studies. Section 3 introduces the preliminary concepts and notation. Section 4 describes the proposed methodology and computational pipeline. Section 5 presents the results. Section 6 discusses the findings, limitations, and future directions. Section 7 concludes the study.

## 2 Related Studies

Signaling-pathway modeling has become an important computational approach for studying cancer-associated dysregulation and therapeutic response. Growth Factor (GF), PI3K/AKT/mTOR, and MAPK signaling networks are particularly important because abnormal activation of these pathways can promote uncontrolled proliferation, survival, metabolic reprogramming, and treatment resistance. Earlier pathway-logic studies modeled the GF pathway using potentially faulty signaling components and investigated how pathway perturbations could lead to abnormal output activation [9]. Similarly, MAPK-centered pathway models have been explored in cancer and metabolic contexts, especially because Ras/RAF/MEK/ERK and PI3K-associated branches are central to growth-factor response and proliferative signaling [4]. Previous studies have also examined multi-node inhibition and drug-repurposing strategies within the MAPK pathway, highlighting the importance of evaluating intervention effects beyond isolated single targets [8].

Dynamic Bayesian Networks have also been used to model biological regulatory systems, especially gene regulatory networks, because they can represent temporal dependencies among biological variables [15]. Recent advances in computational biology have increasingly relied on network-based and omics-driven approaches to investigate disease mechanisms and identify potential biomarkers. For instance, analyses of miRNA-mRNA interaction networks have been used to uncover regulatory patterns associated with disease [16], while studies of melanoma-associated lncRNAs have highlighted the benefits of combining sequence, structural, and interaction information to support disease-related gene characterization [17]. These studies show that network-aware and omics-driven computational approaches can provide useful biological insight across different disease contexts.

In pathway-level therapeutic modeling, Boolean and logic-based models remain common because they provide interpretable qualitative representations when detailed kinetic parameters are unavailable. Boolean modeling has been widely used to study signaling and regulatory networks [18], while logic-based models have been applied to analyze mammalian cell signaling and signal-transduction data [19], [20]. More recent reviews have highlighted the continued use of Boolean and logical modeling approaches in systems medicine and cancer signaling research [14], [21]. These studies show that logical models remain valuable for cancer signaling analysis, especially when mechanistic interpretability is required.

Several prior studies have investigated drug-response prediction and intervention design using pathway logic. Layek et al. studied cancer therapy design using pathway logic and explored drug inhibition points in the GF pathway [9]. Muhuri et al. further investigated drug-vector minimization in cancer therapy using Boolean models of gene regulatory networks [22]. Max-SAT-based strategies have also been applied to identify intervention combinations for specific fault patterns in signaling networks [23]. Multiple-fault drug application in signaling pathways has also been studied using Boolean network-based modeling, supporting the importance of evaluating larger fault spaces rather than only single-fault cases [7]. These works established formal logic and optimization-based methods as useful tools for computational therapy design, although many of them relied on deterministic or discrete-state propagation.

More recent work has extended logical modeling toward personalized and drug-combination analysis. Montagud et al. developed patient-specific Boolean models of signaling networks to guide personalized treatment prioritization [24]. In breast cancer, machine-learning-based subtype prediction has also been investigated using gene expression signatures, showing how omics-driven computational models can support cancer stratification and personalized interpretation [25]. Together, these studies demonstrate the increasing use of pathway-level, omics-driven, and machine-learning models for personalized and combination-therapy analysis.

Stochastic and probabilistic extensions of logical modeling have been proposed to address uncertainty in biological systems. Shmulevich et al. introduced Probabilistic Boolean Networks as a rule-based uncertainty model for gene regulatory networks [26]. Stochastic Boolean simulation environments, such as MaBoSS, further demonstrate that Boolean-like models can produce time-dependent probabilities for genes, proteins, and phenotypes [27]. Similar probabilistic frameworks have also been applied to model biological state transitions and temporal dynamics using time-course data [28]. These developments are relevant because they move beyond purely deterministic state transitions and support probability-based interpretation of biological network dynamics.

Drug-discovery and therapeutic-prioritization studies have also increasingly used machine learning and network-guided strategies. Deep learning-based protein-ligand binding affinity models, such as CGDeepAff and RLBindDeep, illustrate how neural architectures can support molecular interaction prediction and computational drug-discovery pipelines [29], [30]. Network-informed offline reinforcement learning has also been proposed for pharmacogenomic drug prioritization, showing the value of explicitly incorporating biological network structure into therapeutic decision-making [31].

Despite these advances, several gaps remain. Many studies focus on deterministic Boolean propagation, specific patient or cell-line contexts, or predefined drug-target sets. Fewer studies systematically evaluate probabilistic temporal DBN behaviour across increasing multi-fault orders while comparing known-drug combinations, burden-aware Pareto ranking, and custom dual-target intervention hypotheses within a single framework. This study addresses that gap by modeling the GF and MAPK pathways using a probabilistic temporal DBN, evaluating one-to four-fault dysregulation scenarios, ranking known-drug vectors using encoded-burden reduction, and identifying high-impact custom dual-target intervention candidates.

## 3 Preliminaries

### 3.1 Dynamic Bayesian Networks for Signaling Pathways

A Dynamic Bayesian Network (DBN) is a probabilistic graphical model used to represent systems whose variables evolve over discrete time steps [3]. In a DBN, nodes represent random variables, while edges represent probabilistic dependencies among them. For biological signaling pathways, this representation is useful because pathway components do not act independently; upstream receptors, adaptors, kinases, inhibitors, transcription factors, and other regulatory molecules often influence the state of one protein. A signaling pathway can be represented as a directed graph *G* = (*V, E*), where *V* is the set of nodes and *E*⊆ *V* ×*V* is the set of directed regulatory interactions. Each node (*v* ∈*V*) represents a biological entity in the pathway, such as a growth factor, receptor, intracellular signaling protein, transcription factor, or residual output protein. Each directed edge (*u, v*) ∈ *E* indicates that node *u* has a regulatory influence on node *v*.

For any node (v), the set of parent nodes is defined as:

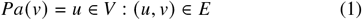

The parent set (*Pa* (*v*)) contains the upstream regulators that directly determine or influence the state of node (*v*). In signaling-pathway modeling, these parent-child relationships correspond to biochemical interactions such as activation, inhibition, complex formation, and convergent signaling.

The node set can be divided into three broad categories:

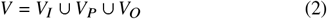

where (*V*_*I*_) denotes input nodes, (*V*_*P*_) denotes internal protein nodes, and (*V*_*O*_) denotes output nodes. Input nodes represent external or upstream growth-factor signals. Internal protein nodes represent signaling intermediates within the pathway. Output nodes represent downstream biological readouts, such as transcription-factor activation or residual signaling effects.

This structure allows the pathway to be interpreted as a signal-propagation system. External signals enter through the input nodes, pass through internal regulatory interactions, and eventually influence the output nodes. In a temporal DBN, this propagation is evaluated over discrete time points, allowing the network state to evolve from an initial input condition toward a final output profile [32], [33].

### 3.2 Probabilistic Protein-State Representation

In pathway-logic models, a protein is often described as either active or inactive. This binary interpretation can be represented using a random variable as:

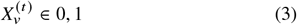

where 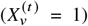 denotes that protein (*v*) is active at time (*t*), and 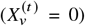denotes that it is inactive or inhibited. However, biological signaling is rarely perfectly binary. Protein activation may be partial, uncertain, noisy, context-dependent, or affected by incomplete inhibition and incomplete fault penetrance. Therefore, instead of propagating only hard binary states, the DBN represents the state of each node through its activation probability:

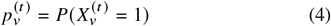

where,

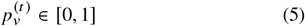

A value close to 1 indicates a high probability of activation, while a value close to 0 indicates a low probability of activation. Intermediate values represent partial or uncertain activation. This probabilistic interpretation is particularly suitable for biological systems, where signaling events may be influenced by stochastic molecular interactions, pathway crosstalk, regulatory feedback, and imperfect therapeutic effects.

For a pathway with (*m*) input nodes, the initial input condition can be represented as:

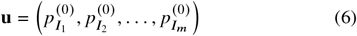

In the considered signaling pathways, the input nodes correspond to growth-factor or upstream signaling stimuli. The input vector may be initialized using binary values to represent the presence or absence of external stimuli, while the downstream nodes evolve as probabilities during temporal DBN propagation.

At each time step, the activation probability of a non-input node depends on the activation probabilities of its parent nodes. Therefore, for (*v* ∈ *V*_*P*_ ∪ *V*_*O*_), the general form of node-state propagation can be written as:

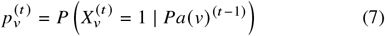

This expression states that the activation probability of node (*v*) at time (*t*) depends on the states of its parent nodes at the previous time step. The exact form of this dependency is determined by the biological interaction rule associated with that node.

### 3.3 Soft-Logic Representation of Biochemical Interactions

Biochemical signaling interactions can often be described using logical rules. For example, a protein may be activated by an upstream protein, inhibited by another protein, activated when either of two parent proteins is active, or activated only when multiple parent proteins are simultaneously active. In a probabilistic DBN, such relationships can be represented using soft-logic rule functions.

For a non-input node (*v*), let ( *f*_*v*_) denote the rule function associated with that node. The rule function maps the activation probabilities of the parent nodes into an ideal activation probability:

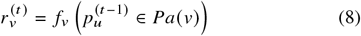

where 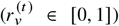 is the rule-derived activation probability before any additional model components, such as persistence, fault effects, or therapeutic interventions, are considered.

The most common soft-logic rules used in pathway modeling are identity, NOT, OR, AND, and NOR, which are shown in Table 1.

**Table 1:** Soft-logic rules for pathway modeling.

| Rule type | Biological/logical interpretation | Probabilistic rule function |
| --- | --- | --- |
| Identity | A parent directly activates the child node. | $f(A) = p_A$ |
| NOT | A parent inhibits the child node. | $f(A) = 1 - p_A$ |
| OR | The child node can be activated by at least one parent. | $f(A_1, \dots, A_m) = 1 - \prod_{i=1}^m (1 - p_{A_i})$ |
| AND | The child node requires simultaneous activation of all parents. | $f(A_1, \dots, A_m) = \prod_{i=1}^m p_{A_i}$ |
| NOR | The child node is active only when none of the parent nodes are active. | $f(A_1, \dots, A_m) = \prod_{i=1}^m (1 - p_{A_i})$ |

The identity rule represents direct activation. If the upstream node has a high activation probability, the downstream node also receives a high activation signal. The NOT rule represents inhibition, where a high upstream activation probability suppresses the downstream node. The OR rule models alternative activation routes, where any one active parent may activate the child node. The AND rule models joint dependency, where all parent nodes must be active to strongly activate the child node. The NOR rule represents a negative convergent relationship, where activation occurs only when all parent signals are absent.

For more complex regulatory cases, a general truth-table representation can be used. Suppose a node has (*m*) parent nodes (*A*_1_, *A*_2_, …, *A*_*m*_). Let,

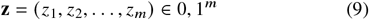

denote a binary configuration of the parent states, and let (*y*_**z**_ ∈0, 1) denote the corresponding output of the rule for that parent configuration. The probabilistic rule output can then be written as:

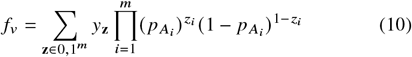

This expression calculates the expected output of a Boolean rule under probabilistic parent activation. When all parent probabilities are exactly 0 or 1, the soft-logic rule gives the corresponding Boolean output. When parent probabilities take intermediate values, the rule gives the expected activation probability over all possible parent-state configurations. Thus, soft-logic functions provide a bridge between discrete pathway logic and probabilistic temporal modeling.

### 3.4 Temporal Propagation in Probabilistic Pathway Models

Temporal propagation allows a pathway model to capture how signaling states evolve over time. In a static model, each node is evaluated once from the input condition. In a temporal DBN, node states are updated across multiple discrete time steps:

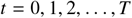

The time horizon (*T*) determines how long the pathway is allowed to evolve. At (*t* = 0), input nodes are initialized according to the input vector, and internal and output nodes are initialized according to the model setup. At each subsequent time step, the activation probability of each non-input node is updated based on the previous-time-step probabilities of its parents.

A general temporal update can be interpreted as a combination of parent-derived signaling and state persistence. Parent-derived signaling reflects the current regulatory influence received from upstream nodes. State persistence reflects the tendency of a node to retain part of its previous activation state. This is useful for representing biological memory, delayed deactivation, or gradual pathway transitions.

In this type of model, the final output vector at time (*T*) is used to summarize the pathway response:

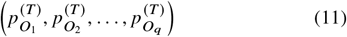

where (*q*) is the number of output nodes. The values in (**o**^(*T*)^) represent the final activation probabilities of the downstream readout nodes. These output probabilities can then be used in downstream analyses, such as fault-response evaluation, intervention assessment, and ranking of candidate drug combinations.

### 3.5 Stuck-at Faults in Signaling Pathways

Faults in signaling pathways represent persistent abnormalities that disrupt normal signal transmission. In biological systems, such abnormalities may arise due to mutation, constitutive activation, loss of function, altered expression, or disrupted regulation. These faults can cause downstream pathway components to become active or inactive even when the corresponding upstream biological condition would not normally produce that state.

A common abstraction for such abnormalities is the stuck-at fault. In a stuck-at fault model, a faulty node is biased toward a fixed state. Two major types of stuck-at faults are commonly considered:

i. **Stuck-at-1 fault:** The node behaves as if it is constitutively active.
ii. **Stuck-at-0 fault:** The node behaves as if it is persistently inactive or inhibited.

A stuck-at-1 fault can represent gain-of-function behavior, constitutive signaling activation, or abnormal oncogenic activation. A stuck-at-0 fault can represent loss-of-function behavior, persistent suppression, or defective signaling transmission. In cancer-related pathway models, stuck-at-1 faults are particularly relevant for proteins whose abnormal activation can drive proliferation, while stuck-at-0 faults are relevant for proteins whose suppression removes inhibitory control or disrupts apoptotic regulation.

For a faulty node *f*, a deterministic stuck-at-1 condition can be represented as

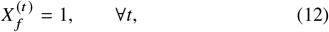

whereas a deterministic stuck-at-0 condition can be represented as

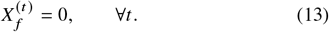

In a probabilistic DBN setting, the same idea can be implemented by replacing the normal conditional probability distribution of the faulty node with a fault-biased distribution. For example, with a small tolerance parameter *ϵ*, the stuck-at-1 and stuck-at-0 cases may be written as

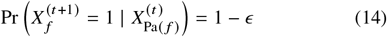

and

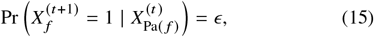

respectively. The deterministic case is obtained when *ϵ* = 0.

For a pathway with a set of fault-candidate nodes *C* ⊆ *P*, a concurrent *k*-fault scenario is defined by selecting a subset of *k* faulty nodes,

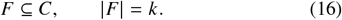

If both stuck-at-0 and stuck-at-1 conditions are allowed, then a complete fault scenario is represented as

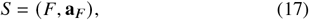

where **a**_*F*_ ∈{0, 1 }^| *F* |^ specifies the stuck-at state assigned to each node in *F*. The complete set of concurrent *k*-fault scenarios is therefore

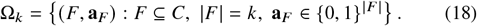

Thus, when both stuck-at states are considered, the total number of possible *k*-fault scenarios is

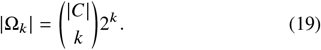

This combinatorial growth motivates the need for efficient computational evaluation, particularly when multiple simultaneous faults are examined across large pathway models.

Fig. 1 and Fig. 2 represent the corresponding DBN for GF and MAPK pathways, respectively. Each protein in the pathway is assigned a numeric identifier denoting the protein’s location and possible fault location.

**Fig. 1:**
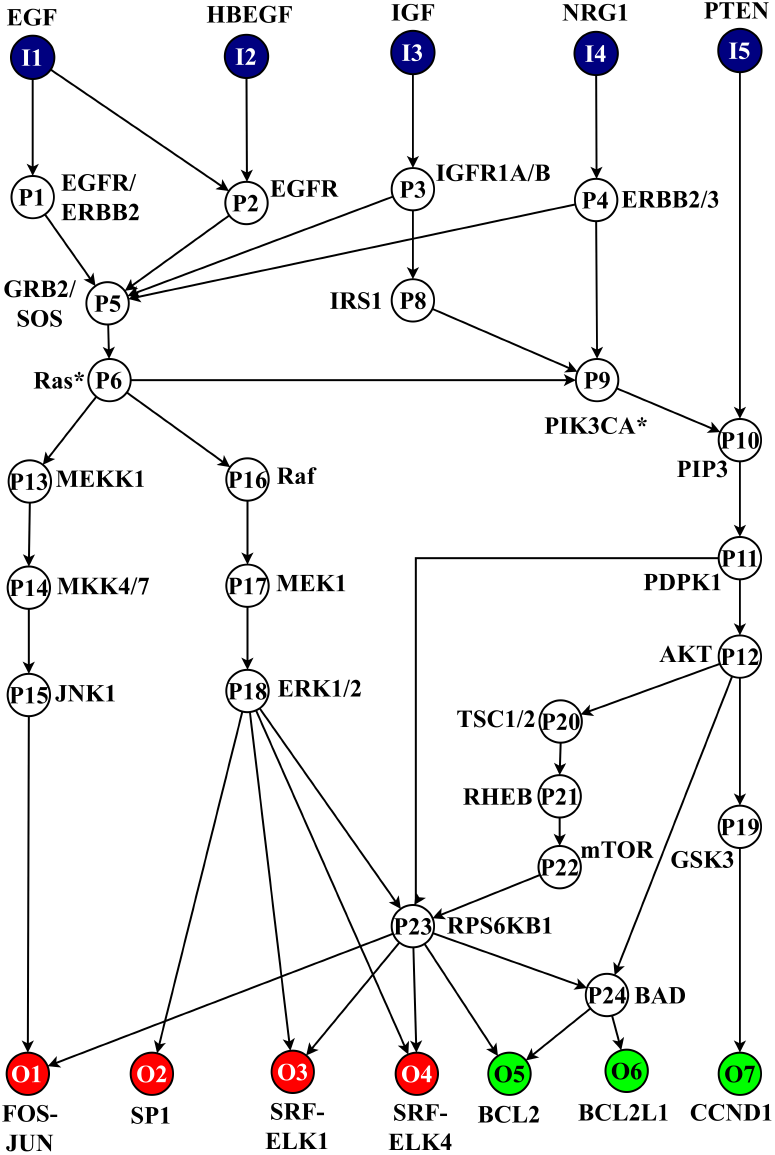
Dynamic Bayesian network representation of the growth factor signaling pathway. Blue nodes denote external input signals, white nodes denote internal pathway components, and terminal colored nodes denote downstream output readouts. Each node is assigned a unique identifier to support systematic stuck-at fault injection and downstream effect analysis.

**Fig. 2:**
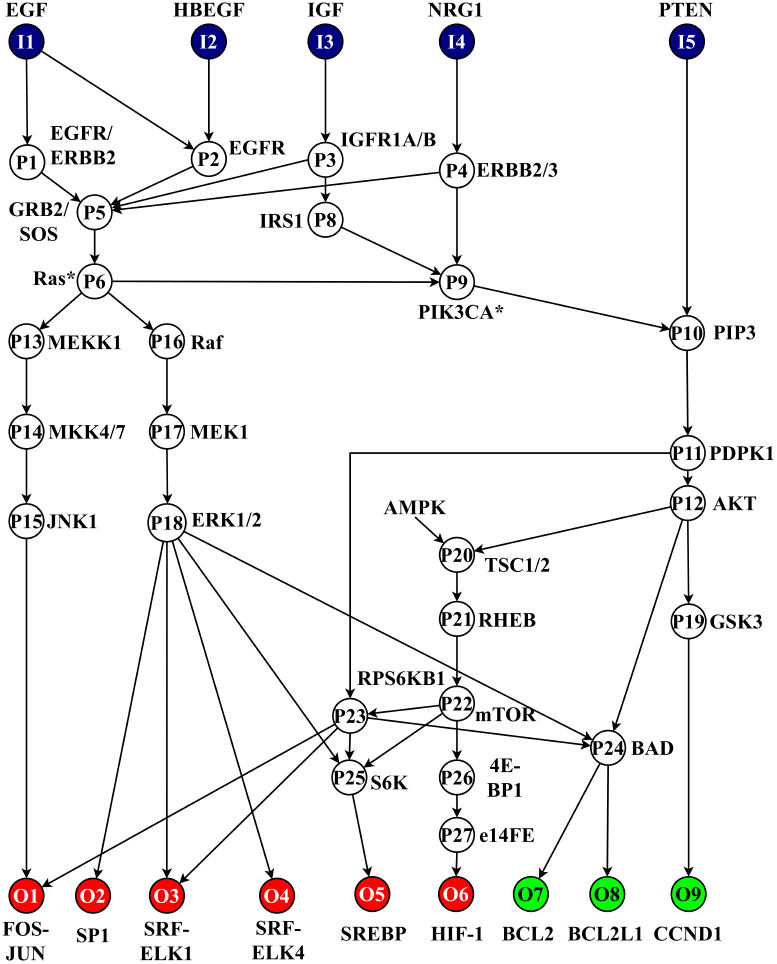
Dynamic Bayesian network representation of the MAPK-associated signaling pathway. The model captures receptor-level activation, Ras/MAPK signaling, PI3K/AKT/mTOR signaling, AMPK/TSC/RHEB regulation, and multiple downstream transcriptional, survival, metabolic, and proliferative outputs. Node identifiers are used to define fault-candidate locations during stuck-at fault simulation.

### 3.6 Drug Interventions in Signaling Pathway Models

Drug interventions in signaling-pathway models represent therapeutic suppression or modulation of selected pathway components. In cancer-associated networks, abnormal activation of receptors, kinases, and downstream proteins can sustain proliferative signaling even without normal extracellular stimulation. Targeted drugs may reduce this activity by inhibiting specific proteins or activating regulatory components that indirectly suppress pathway output. Thus, pathway-level models represent each drug according to its known biological point of action, such as receptor inhibition, kinase blockade, or indirect downstream regulatory control, and these drug-target relationships form the basis for constructing candidate drug combinations in the computational framework.

The present study considers seven known drugs that have been used in prior signaling-pathway intervention analyses: Lapatinib, AG825, AG1024, U0126, LY294002, Temsirolimus, and Metformin. The pathway-level inhibition or modulation points associated with these drugs are summarized in Table 2.

**Table 2:** Drugs and the corresponding proteins inhibited.

| Index | Drug | Proteins suppressed within each signaling network |  |
| --- | --- | --- | --- |
|  |  | GF | MAPK |
| 1 | LAPATINIB | EGFR/ERBB2 | EGFR/ERBB2 |
|  |  | EFGR | EFGR |
|  |  | ERBB2/3 | ERBB2/3 |
| 2 | AG825 | EGFR/ERBB2 | EGFR/ERBB2 |
|  |  | ERBB2/3 | ERBB2/3 |
| 3 | AG1024 | IGFR1A/B | IGFR1A/B |
| 4 | U0126 | MEK1 | MEK1 |
| 5 | LY294002 | PIK3CA | PIK3CA |
| 6 | TEMSIROLIMUS | mTOR | mTOR |
| 7 | METFORMIN | TCS1/2 | AMPK |

The first six drugs are represented as inhibitory interventions on their corresponding pathway targets. Lapatinib and AG825 act on the EGFR/ERBB receptor family, AG1024 targets IGF receptor signaling, U0126 inhibits MEK1, LY294002 inhibits PI3K/PIK3CA signaling, and Temsirolimus targets mTOR. These interventions are relevant because receptor tyrosine kinase signaling, PI3K/AKT/mTOR signaling, and Ras/MEK/ERK signaling are frequently dysregulated in cancer-associated pathway activation.

Metformin is treated differently from the direct kinase and receptor inhibitors because its pathway-level effect is commonly linked to AMPK activation. AMPK activation can suppress mTOR-associated signaling through regulatory intermediates such as TSC1/2. Therefore, in pathway-level intervention modeling, Metformin is represented as a regulatory intervention associated with TSC1/2 or AMPK-mediated control rather than as a direct inhibitor of the same type as the kinase-targeting, a phenomenon known as the “Warburg effect” [34].

Table 2 defines the biological drug-target knowledge used as input to the computational framework. The exact mathematical representation of drug action, including how drug targets are mapped to pathway node identifiers and how inhibitory or activating interventions modify node activation probabilities, is described in the methodology section.

### 3.7 Pareto Dominance and Multi-objective Ranking

[35] Therapeutic candidate ranking often involves more than one objective. For example, one may want to maximize pathway-correction efficacy while minimizing the number of drugs, targets, or intervention points. These objectives can conflict with each other: a candidate with high efficacy may require many drugs, while a simpler candidate may have lower efficacy. Pareto dominance provides a formal way to compare candidates under such competing objectives.

Consider two candidate interventions, (*a*) and (*b*). Suppose each candidate is evaluated using two objective functions: one objective to be maximized, such as efficacy, and one objective to be minimized, such as intervention burden. Candidate (*a*) is said to dominate candidate (*b*) if (*a*) is at least as good as (*b*) in all objectives and strictly better in at least one objective.

For a maximization objective (*M* (*c*)) and a minimization objective (*B*(*c*)), candidate (*a*) dominates candidate (*b*) if

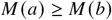

And

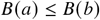

with at least one strict inequality. The Pareto front is the set of candidates that are not dominated by any other candidate:

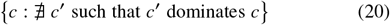

Candidates on the Pareto front represent trade-off solutions. No candidate on the front can be improved in one objective without worsening at least one other objective. In the context of signaling-pathway intervention analysis, Pareto dominance is useful for identifying candidate drug combinations or target combinations that balance pathway-level efficacy with intervention simplicity.

### 3.8 Summary of Notation

The major notation used throughout the manuscript is summarized in Table 3.

**Table 3:** Notation details used in the study.

| Symbol | Meaning |
| --- | --- |
| $G = (V, E)$ | Directed graph representation of a signaling pathway |
| $V$ | Set of all pathway nodes |
| $E$ | Set of directed regulatory interactions |
| $V_I$ | Set of input or growth-factor nodes |
| $V_P$ | Set of internal protein nodes |
| $V_O$ | Set of output nodes |
| $Pa(v)$ | Parent set of node $(v)$ |
| $X_v^{(t)}$ | Binary random variable representing the state of node $(v)$ at time $(t)$ |
| $p_v^{(t)}$ | Activation probability of node $(v)$ at time $(t)$ |
| $\mathbf{u}$ | Input vector supplied to the pathway |
| $f_v$ | Soft-logic rule function associated with node $(v)$ |
| $r_v^{(t)}$ | Rule-derived activation probability of node $(v)$ at time $(t)$ |
| $T$ | Fixed temporal simulation horizon |
| $\mathbf{o}^{(T)}$ | Output-probability vector at the final time horizon |
| $C$ | Set of fault-candidate nodes |
| $F$ | Specific set of faulty nodes |
| $k$ | Number of concurrent faults |
| $\mathcal{F}_k$ | Set of all $k$-fault combinations |
| $\mathbf{d}$ | Binary drug-combination vector |
| $\mathcal{P}$ | Pareto front |

## 4 Proposed Methodology

The proposed framework models each signaling pathway as a probabilistic temporal Dynamic Bayesian Network (DBN). Each protein or pathway component is represented as a node, and each directed biochemical interaction is represented as a probabilistic dependency. Instead of propagating only binary active/inactive states, the framework propagates activation probabilities over a fixed number of discrete time steps.

For each pathway, the model accepts an input vector representing the activation status of growth-factor input nodes. The network then evolves through soft-logic update rules, fault perturbations, and therapeutic interventions. The final output probabilities are compressed into an encoded pathway-burden score, which is used to compare fault-only and intervention-treated states. Known drug combinations and custom dual-target interventions are then ranked according to their ability to reduce the encoded burden relative to a no-intervention baseline.

### 4.1 Pathway Graph Construction and Node Mapping

Each pathway is represented as a directed acyclic graph *G* = (*V, E*), where *V* is the set of nodes and *E* is the set of directed interactions. The node set is divided into input nodes, protein nodes, and output nodes, *V* = *V*_*I*_ ∪ *V*_*P*_∪ *V*_*O*_.

Both pathways use five input nodes, (*I*_1_) to (*I*_5_), representing upstream growth-factor or pathway-entry signals. The internal protein nodes are denoted as (*P*_*i*_), and the output nodes are denoted as (*O*_*i*_). The output nodes are further divided into transcription-factor outputs and residual-protein outputs.

The pathway-specific node composition used in the model is summarized in Table 4.

**Table 4:**
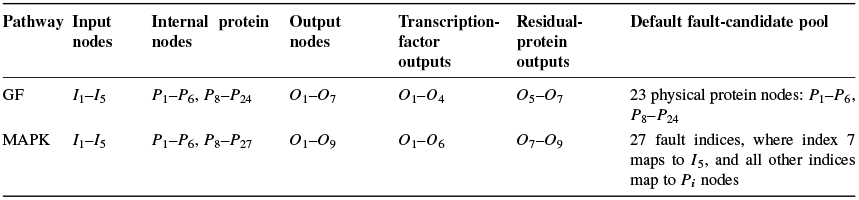
Pathway-specific node sets and default fault-candidate pools.

| Pathway | Input nodes | Internal protein nodes | Output nodes | Transcription-factor outputs | Residual-protein outputs | Default fault-candidate pool |
| --- | --- | --- | --- | --- | --- | --- |
| GF | $I_1-I_5$ | $P_1-P_6, P_8-P_{24}$ | $O_1-O_7$ | $O_1-O_4$ | $O_5-O_7$ | 23 physical protein nodes: $P_1-P_6, P_8-P_{24}$ |
| MAPK | $I_1-I_5$ | $P_1-P_6, P_8-P_{27}$ | $O_1-O_9$ | $O_1-O_6$ | $O_7-O_9$ | 27 fault indices, where index 7 maps to $I_5$, and all other indices map to $P_i$ nodes |

For the GF pathway, no physical (*P*_7_), (*P*_25_), (*P*_26_), or (*P*_27_) node is included in the implemented graph. Therefore, the default GF fault pool contains 23 physical protein nodes. For the MAPK pathway, index 7 is treated as an input-level fault corresponding to (*I*_5_), while all other indices correspond to protein nodes.

The rule structure of the core GF and MAPK pathway is summarized in Table 5. These rules are evaluated using the probabilistic soft-logic functions described in Section 3.3.

**Table 5:** Core pathway rule structure shared by the GF and MAPK DBN models.

| Node | Rule type | Parent nodes | Interpretation |
| --- | --- | --- | --- |
| $P_1$ | Identity | $I_1$ | Direct activation |
| $P_2$ | OR | $I_1, I_2$ | Alternative activation |
| $P_3$ | Identity | $I_3$ | Direct activation |
| $P_4$ | Identity | $I_4$ | Direct activation |
| $P_5$ | OR | $P_1, P_2, P_3, P_4$ | Convergent receptor/adaptor signaling |
| $P_6$ | Identity | $P_5$ | Direct propagation |
| $P_8$ | Identity | $P_3$ | Direct propagation |
| $P_9$ | OR | $P_4, P_6, P_8$ | PI3K-associated convergent activation |
| $P_{10}$ | Truth-table rule | $I_5, P_9$ | Equivalent to $P_{10} = \text{OR}(\text{NOT}(I_5), P_9)$ |
| $P_{11}$ | Identity | $P_{10}$ | Direct propagation |
| $P_{12}$ | Identity | $P_{11}$ | Direct propagation |
| $P_{13}$ | Identity | $P_6$ | MAPK-branch propagation |
| $P_{14}$ | Identity | $P_{13}$ | MAPK-branch propagation |
| $P_{15}$ | Identity | $P_{14}$ | MAPK-branch propagation |
| $P_{16}$ | Identity | $P_6$ | RAF/MEK/ERK branch propagation |
| $P_{17}$ | Identity | $P_{16}$ | MEK-associated propagation |
| $P_{18}$ | Identity | $P_{17}$ | ERK-associated propagation |
| $P_{19}$ | NOT | $P_{12}$ | Inhibitory regulation |
| $P_{20}$ | NOT | $P_{12}$ | TSC1/2- or AMPK-linked regulatory node |
| $P_{21}$ | NOT | $P_{20}$ | Inhibitory regulatory propagation |
| $P_{22}$ | Identity | $P_{21}$ | mTOR-associated propagation |
| $P_{23}$ | OR | $P_{11}, P_{18}, P_{22}$ | Convergent downstream proliferative signaling |
| $P_{24}$ | NOR | $P_{12}, P_{23}$ | Negative convergent regulation |

The GF pathway terminates after (*P*_24_) and contains seven output nodes. The MAPK pathway includes three additional internal nodes (*P*_25_), (*P*_26_), and (*P*_27_), followed by nine output nodes. The additional MAPK-specific rules are summarized in Table 6 and the overall pathway output rules are summarized in Table 7.

**Table 6:** MAPK-specific internal node rules.

| Node | Rule type | Parent nodes |
| --- | --- | --- |
| $P_{25}$ | OR | $P_{18}, P_{22}$ |
| $P_{26}$ | NOR | $P_{18}, P_{22}$ |
| $P_{27}$ | NOT | $P_{26}$ |

**Table 7:**
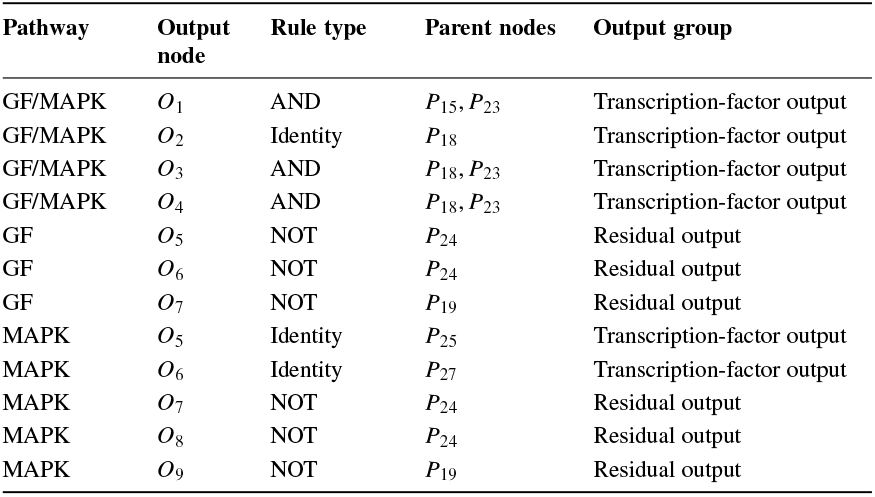
Output-node rule structure and output grouping.

| Pathway | Output node | Rule type | Parent nodes | Output group |
| --- | --- | --- | --- | --- |
| GF/MAPK | $O_1$ | AND | $P_{15}, P_{23}$ | Transcription-factor output |
| GF/MAPK | $O_2$ | Identity | $P_{18}$ | Transcription-factor output |
| GF/MAPK | $O_3$ | AND | $P_{18}, P_{23}$ | Transcription-factor output |
| GF/MAPK | $O_4$ | AND | $P_{18}, P_{23}$ | Transcription-factor output |
| GF | $O_5$ | NOT | $P_{24}$ | Residual output |
| GF | $O_6$ | NOT | $P_{24}$ | Residual output |
| GF | $O_7$ | NOT | $P_{19}$ | Residual output |
| MAPK | $O_5$ | Identity | $P_{25}$ | Transcription-factor output |
| MAPK | $O_6$ | Identity | $P_{27}$ | Transcription-factor output |
| MAPK | $O_7$ | NOT | $P_{24}$ | Residual output |
| MAPK | $O_8$ | NOT | $P_{24}$ | Residual output |
| MAPK | $O_9$ | NOT | $P_{19}$ | Residual output |

### 4.2 Temporal Probabilistic DBN Simulation

Let 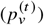 denote the activation probability of node (*v*) at time step (*t*). For each non-input node (*v*), the model first evaluates the ideal soft-logic rule probability:

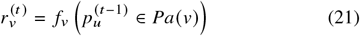

The ideal rule probability is then transformed using the maximum propagation strength (*α*) and basal activation leakage (*η*):

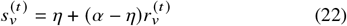

The parameter (*α*) controls the maximum activation probability reached when the ideal rule is fully active. The parameter (*η*) represents the basal activation probability retained even when the ideal rule is inactive.

Temporal persistence is incorporated through (*λ*), which combines the previous activation probability with the current softened rule probability:

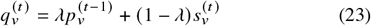

where 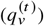 denotes the pre-fault and pre-intervention probability of node (*v*) at time (*t*). Larger values of (*λ*) create stronger temporal memory, whereas (*λ* = 0) produces an instantaneous parent-driven update.

For input nodes, the values are held fixed according to the selected input vector unless the node is explicitly affected by an input-level fault. For non-input nodes, the temporal update is performed synchronously: all nodes at time (*t*) are computed from the state at time (*t* − 1), followed by fault and intervention effects. The output convergence difference at time (*t*) is calculated as:

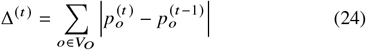

A simulation is recorded as converged if:

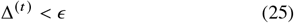

However, all main analyses use the fixed final horizon (*T*) to maintain consistent comparison across fault and intervention scenarios. The fixed-horizon probabilistic temporal DBN simulation is summarized in Algorithm 1.

#### Algorithm 1

Fixed-horizon probabilistic temporal DBN

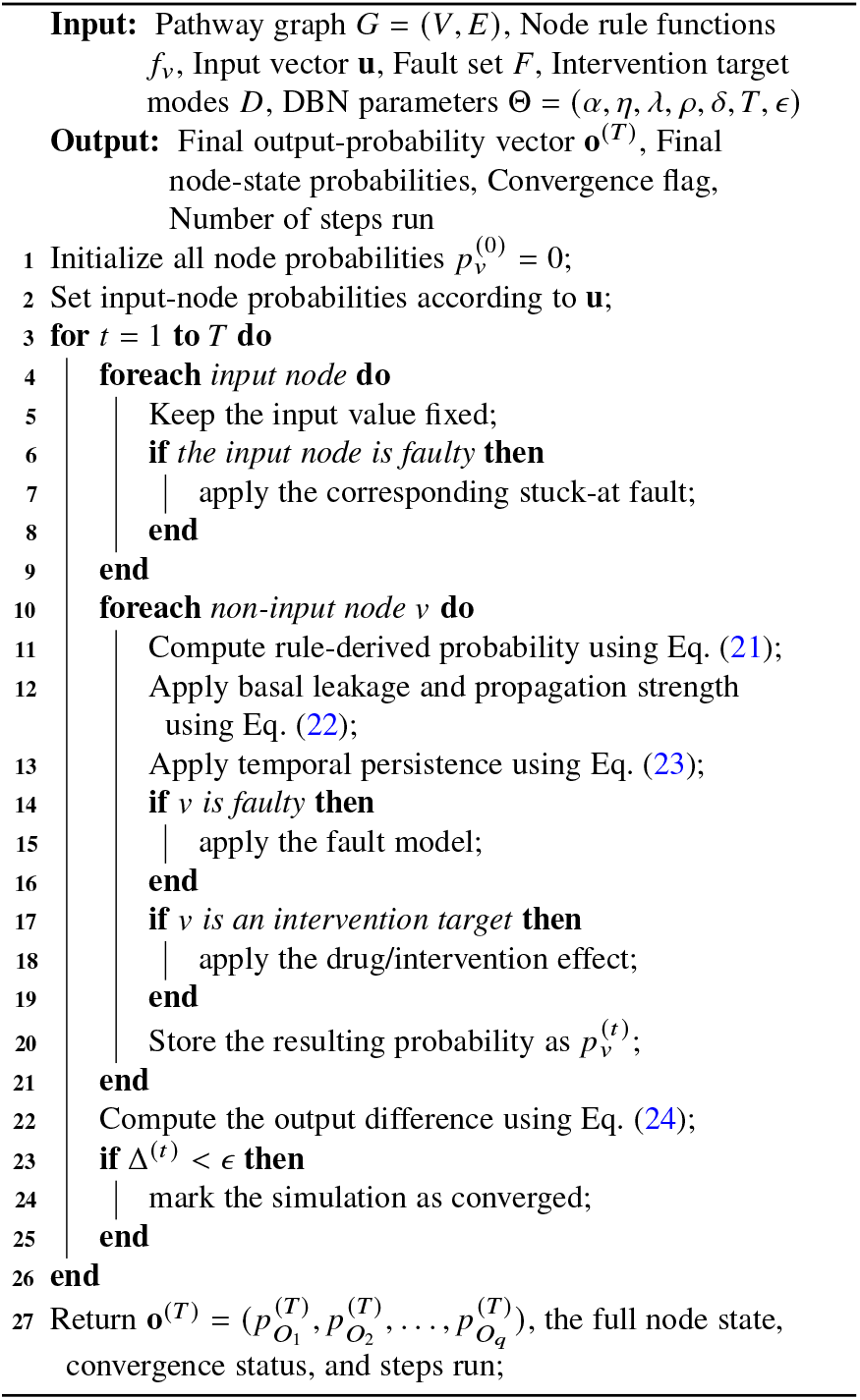

### 4.3 Unique Input Vector Selection

Before fault and drug simulations, the model identifies the input vector that produces the lowest downstream output activation under normal, fault-free conditions. Since both the GF and MAPK pathways have five input nodes, there are 2^5^ = 32 possible binary input vectors.

Let = *u* {0, 1} ^5^ be the set of all possible input vectors. For each input vector (**u** ∈*u*), the fault-free and intervention-free DBN is simulated until the fixed horizon (*T*). The total output burden for (**u**) is calculated as:

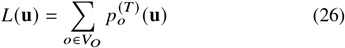

The selected input vector is the one that minimizes the final output burden:

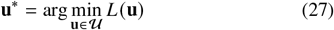

The selected input vector is then used consistently across all baseline, fault, known-drug, custom-target, Pareto, sensitivity, and trajectory simulations. Algorithm 2 describes the unique input selection procedure.

#### Algorithm 2

Unique input vector selection

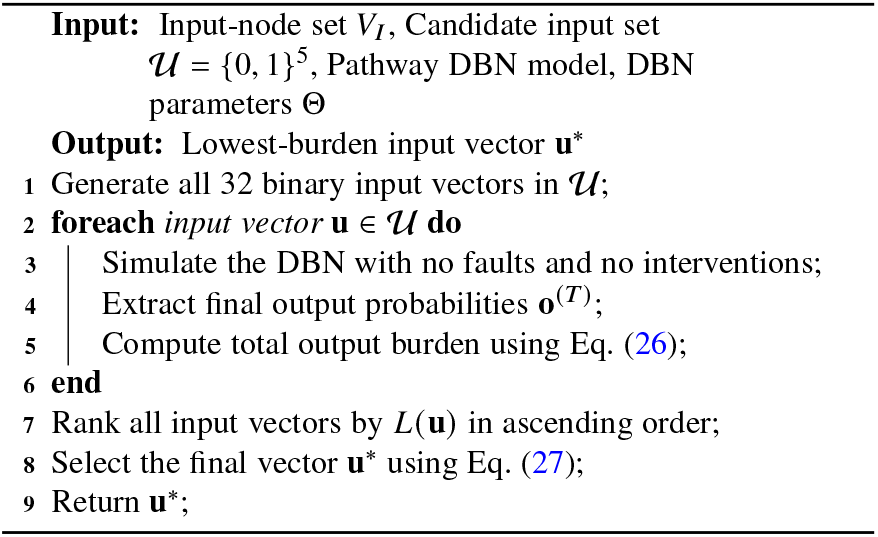

### 4.4 Multi-fault Modeling and Fault Enumeration

Faults are introduced into the DBN to simulate persistent dysregulation of pathway components. A fault-candidate pool (*C*) is defined for each pathway. For a fault order *k*, each concurrent fault scenario is represented as a subset:

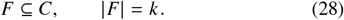

The set of all *k*-fault combinations is defined as:

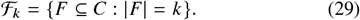

The number of possible *k*-fault combinations is:

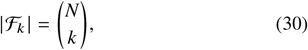

where *N* = |*C*| is the number of fault-candidate locations.

Faults are modeled using a penetrance parameter (*ρ*). If node *v* is assigned a stuck-at-1 fault, its activation probability is driven toward *ρ*, whereas if assigned a stuck-at-0 fault, its activation probability is driven toward 1− *ρ*. Therefore, after temporal propagation, the post-fault probability is defined as:

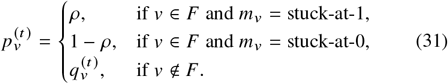

where *m*_*v*_ denotes the fault mode assigned to node *v*, and 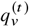is the pre-fault probability obtained after soft-logic propagation and temporal persistence.

The pathway-specific stuck-at-0 nodes are summarized in Table 8. All other default fault candidates are treated as stuck-at-1 nodes. Overall, the fault-only simulation procedure is summarised in Algorithm 3.

**Table 8:** Pathway-specific stuck-at fault modes.

| Pathway | Fault-candidate count | Stuck-at-0 candidates | Stuck-at-1 candidates |
| --- | --- | --- | --- |
| GF | 23 | $P_{19}, P_{20}, P_{24}$ | All remaining GF fault candidates |
| MAPK | 27 | $I_5, P_{19}, P_{20}, P_{24}, P_{26}$ | All remaining MAPK fault candidates |

#### Algorithm 3

Multi-fault baseline simulation

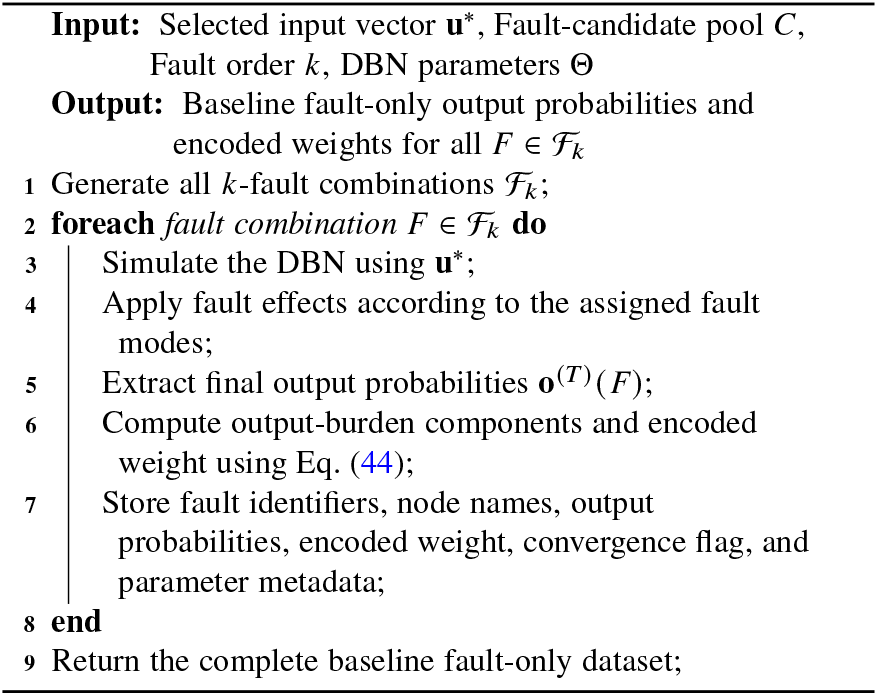

### 4.5 Known-drug Intervention Modeling

The known-drug analysis evaluates all possible combinations of the seven drugs listed in Table 2 under Section 3.6. Each known-drug candidate is encoded as a binary vector:

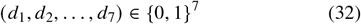

where (*d* _*j*_ = 1) indicates that the *j* -th drug is included in the candidate combination. Since there are seven drugs, the number of candidate drug vectors is 2^7^ = 128. The all-zero vector **d**_0_ = (0, 0, 0, 0, 0, 0, 0) is the no-drug baseline.

The biological drug-target information presented in the preliminaries is mapped to code-level pathway nodes as shown in Table 9.

**Table 9:** Code-level representation of known-drug targets.

| Drug-vector bit | Drug | Biological target interpretation | Code-level DBN target nodes | Main model effect |
| --- | --- | --- | --- | --- |
| 1 | Lapatinib | EGFR/ERBB receptor-family inhibition | $P_1, P_2, P_4$ | Inhibition |
| 2 | AG825 | EGFR/ERBB inhibition | $P_1, P_4$ | Inhibition |
| 3 | AG1024 | IGFR1A/B inhibition | $P_3$ | Inhibition |
| 4 | U0126 | MEK1 inhibition | $P_{17}$ | Inhibition |
| 5 | LY294002 | PIK3CA inhibition | $P_9$ | Inhibition |
| 6 | Temsirolimus | mTOR inhibition | $P_{22}$ | Inhibition |
| 7 | Metformin | AMPK-linked regulatory modulation | $P_{20}$ | AMPK-mode activation |

For an ordinary inhibitory drug target, the intervention effect is modeled as residual suppression:

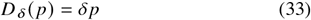

where *p* is the post-fault activation probability of the target node and *δ* ∈[0, 1] is the residual activity parameter. A lower value of *δ* corresponds to stronger inhibition.

For Metformin, the main analysis uses AMPK-mode activation of *P*_20_. This is represented as:

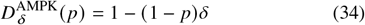

This transformation moves the target node toward a higher activation probability, reflecting the regulatory role of AMPK-linked pathway control. Thus, for a node *v*, the post-intervention probability is defined as:

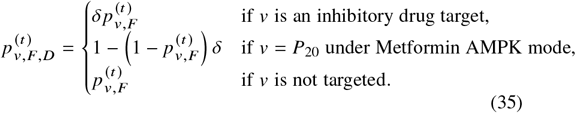

When multiple selected drugs share a target node, the node is treated as a unique intervention target. However, the known-drug intervention burden is counted as the number of active drugs in the drug vector:

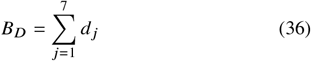

The total number of known drug simulation conditions for fault order *k* is 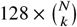. Algorithm 4 summarizes the overall known-drug simulation procedure.

#### Algorithm 4

Known-drug intervention simulation

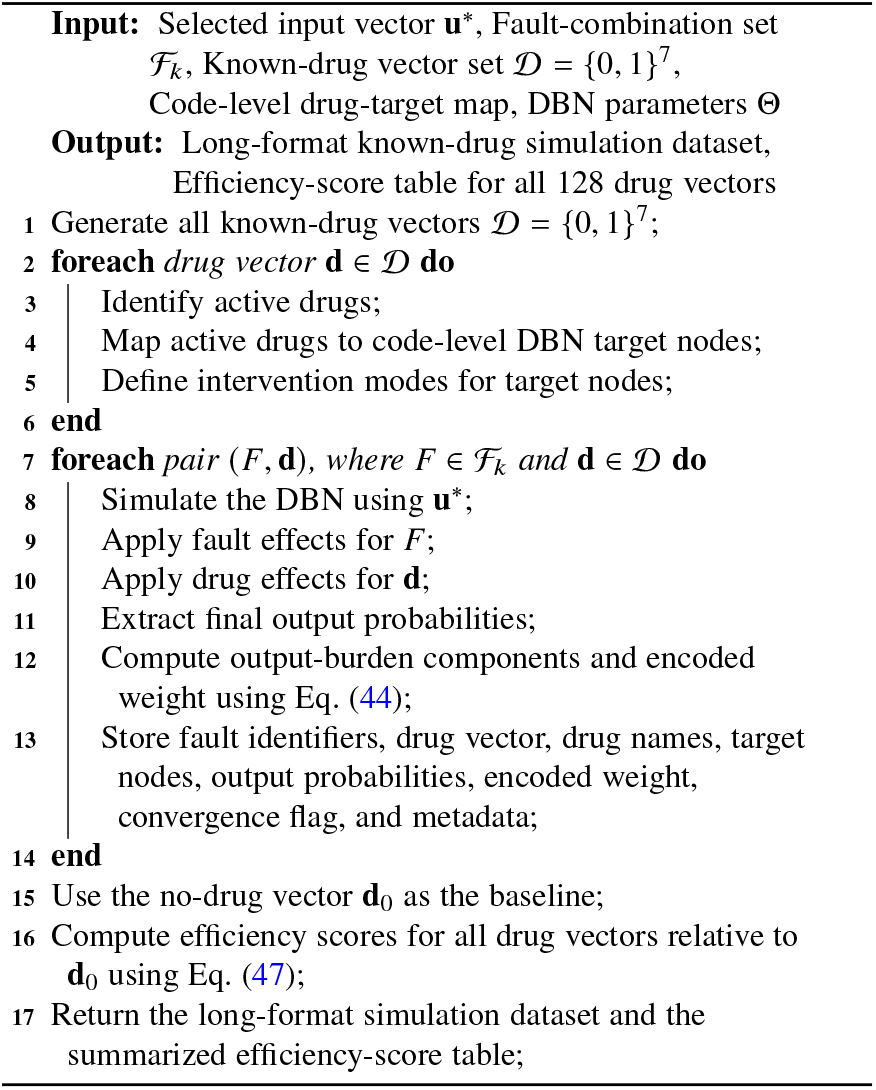

### 4.6 Custom Dual-target Intervention Search

In addition to known-drug combinations, the framework evaluates custom dual-target interventions. These interventions are not interpreted as validated drugs. Rather, they are computational target hypotheses that identify pairs of pathway nodes whose simultaneous intervention may reduce fault-induced output burden.

A custom intervention candidate is represented as:

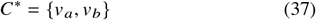

where *v*_*a*_ and *v*_*b*_ are two distinct nodes from the pathway-specific target pool.

For the GF pathway, the default custom target pool is the same set of 23 physical protein fault candidates. Therefore, the number of dual-target candidates is 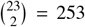. For the MAPK pathway, the default custom target pool contains 26 protein-level targets, excluding the input-level index 7 mapped to *I*_5_. Therefore, the number of default MAPK dual-target candidates is 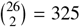.

For efficiency calculation, an empty custom intervention is also included as the baseline candidate:

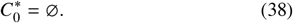

Thus, the total number of custom candidates is 254 for GF and 326 for MAPK under the default setting.

Each selected target in a custom dual-target candidate is treated as an inhibitory target. Therefore, for a node *v*, the post-custom-intervention probability is defined as:

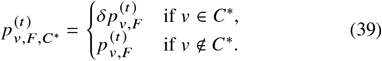

where 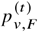 is the post-fault activation probability of node *v* before applying the custom intervention, and *δ* is the residual activity parameter.

The intervention burden for a custom intervention is:

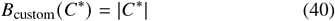

For all non-baseline custom dual-target candidates:

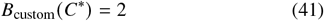

Algorithm 5 summarizes the custom dual-target search.

### 4.7 Encoded Pathway-Burden and Efficiency-score Calculation

After each DBN simulation, the final output probabilities are converted into a scalar encoded pathway-burden value. This encoded burden is then used to compare fault-only and intervention-treated pathway states. The efficiency score quantifies the percentage reduction in encoded burden achieved by a candidate intervention relative to the corresponding no-intervention baseline.

#### 4.7.1 Output Burden and Encoded Weight

After each simulation, the final output-probability vector **o**^(*T*)^ is extracted at the terminal time point *T* :

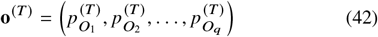

where *q* = 7 for GF and *q* = 9 for MAPK.

The output nodes are divided into transcription-factor outputs and residual-protein outputs. Let *O*_TF_ denote the set of transcription-factor output nodes and *O*_Res_ denote the set of residual-protein output nodes. The corresponding output burdens are defined as:

##### Algorithm 5

Custom dual-target intervention search

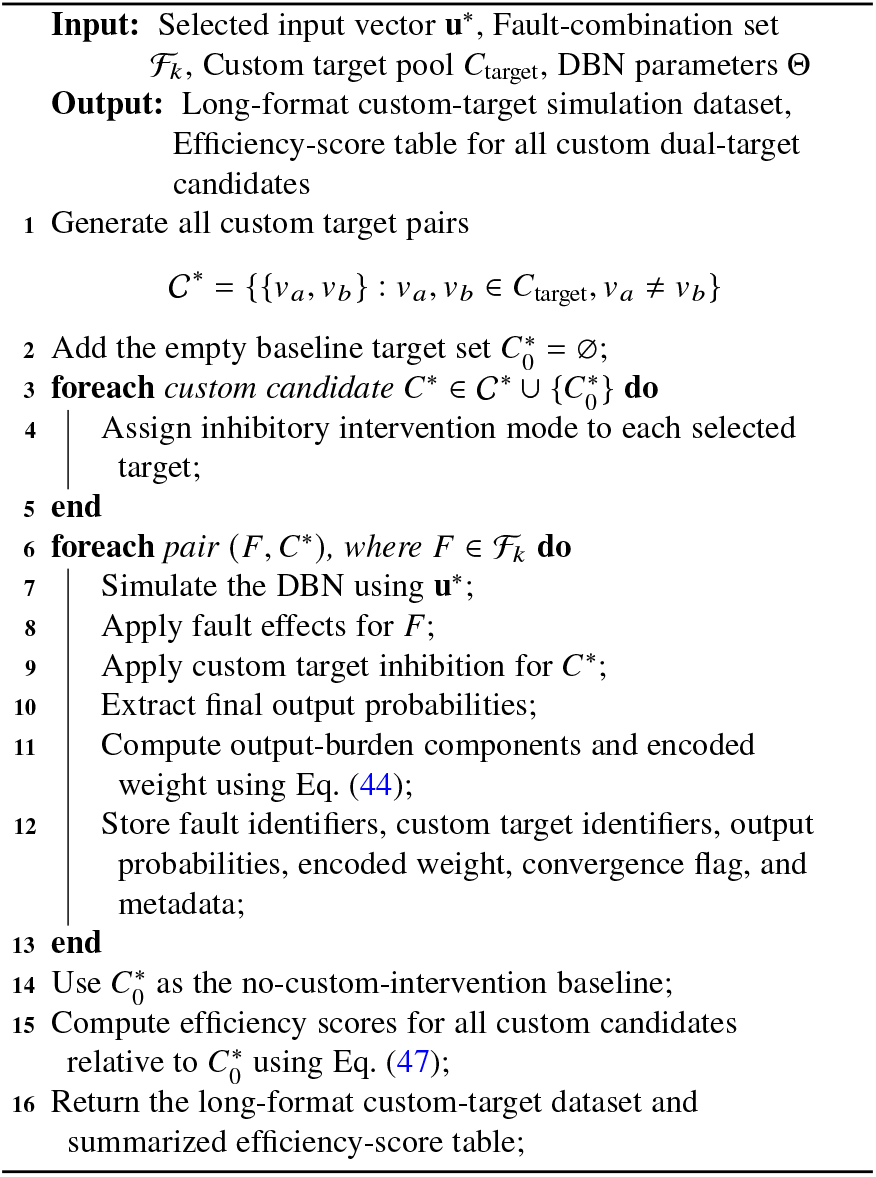

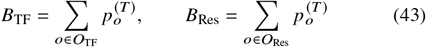

For the GF pathway, the output groups are defined as:

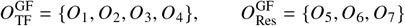

For the MAPK pathway, the output groups are defined as:

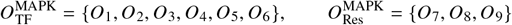

The encoded weight EW compresses the final output-probability vector into a scalar pathway-burden score:

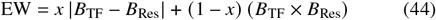

The design parameter *x*∈ (0, 1) controls the relative contribution of burden imbalance and joint burden elevation. The first term penalizes imbalance between transcription-factor and residual-protein activation, while the second term penalizes simultaneous elevation of both output groups. The main analysis uses *x* = 0.5, and the sensitivity analysis evaluates whether candidate rankings remain stable across alternative values of *x*.

#### 4.7.2 Efficiency-score Calculation

For a fixed fault order *k*, each intervention candidate is evaluated across all fault combinations in ( *f*_*k*_). Let *c* denote an intervention candidate. For known-drug analysis, *c* is a drug vector (**d**). For custom-target analysis, *c* is a custom target set (*C*^*\**^).

Let *c*_0_ denote the corresponding no-intervention baseline. For known-drug analysis, the baseline candidate is:

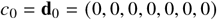

For custom-target analysis, the baseline candidate is:

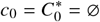

For a fixed fault order *k*, the total encoded burden of the no-intervention baseline is first computed as:

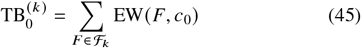

where *c*_0_ denotes the corresponding no-intervention baseline candidate. Similarly, the total encoded burden of an intervention candidate *c* is computed as:

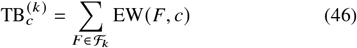

The efficiency score of candidate *c* is then defined as the percentage reduction in total encoded burden relative to the no-intervention baseline:

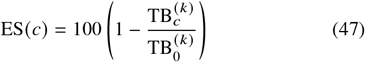

where, EW (*F, c*) denotes the encoded weight obtained when candidate *c* is applied under fault combination *F* using Eq. (44).

A higher value of ES (*c*) indicates a stronger reduction in encoded pathway burden. A value close to zero indicates little improvement over the no-intervention baseline. A negative value indicates that the candidate increases encoded burden relative to the baseline.

For each candidate, the mean encoded weight is also recorded:

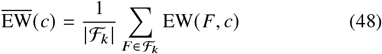

This provides a direct summary of the average final burden produced by the candidate across all fault combinations of the selected order. Algorithm 6 summarizes the encoded-weight and efficiency-score calculation.

### 4.8 Pareto-based Candidate Prioritization

The efficiency score measures the degree of burden reduction, but it does not alone account for the complexity of the intervention. Therefore, the final ranking considers two objectives: maximizing the efficiency score and minimizing the intervention burden:

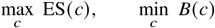

For known-drug candidates, where *c* = **d** and **d** = (*d*_1_, *d*_2_, …, *d*_7_), the intervention burden is defined as the number of active drugs in the drug vector:

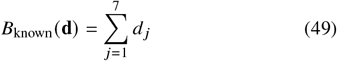

For custom-target candidates, where *c* = *C*^*\**^, the intervention burden is defined as the number of selected target nodes:

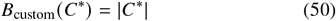

Candidate *a* is said to dominate candidate *b* if *a* is at least as efficient as *b* and no more complex than *b*, with at least one strict improvement:

#### Algorithm 6

Encoded-burden and efficiency-score calculation

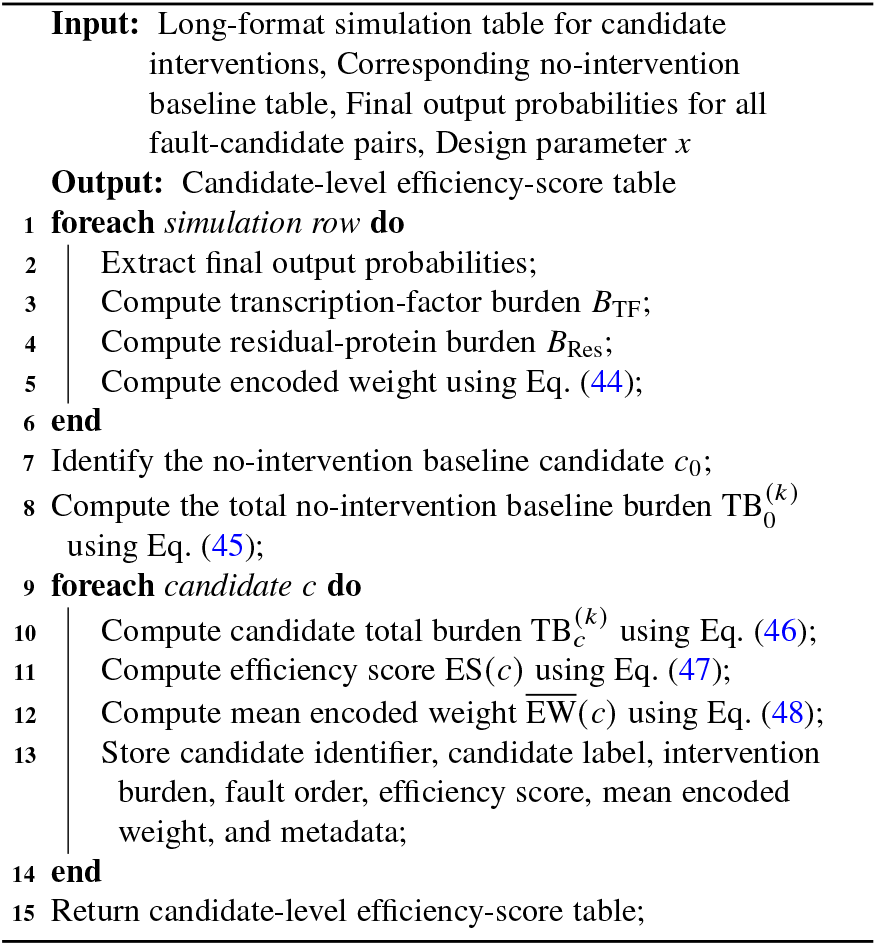

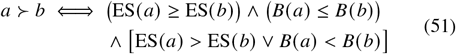

The Pareto front is then defined as the set of candidates that are not dominated by any other candidate:

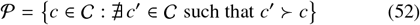

Where *C* denotes the candidate set being evaluated.

Candidates on the Pareto front represent non-dominated tradeoffs between therapeutic efficiency and intervention simplicity.

For known-drug analysis, an additional low-burden candidate subset is extracted as:

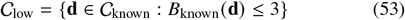

This subset is used to prioritize clinically more interpretable combinations involving one to three drugs. For custom dual-target candidates, all non-baseline candidates have a burden of two by construction. Algorithm 7 summarizes the Pareto-ranking procedure.

### 4.9 Implementation and Simulation Details

The proposed framework requires the repeated simulation of large numbers of fault-intervention conditions. Therefore, the implementation combines batched vectorized execution with a fixed set of default DBN parameters. The batched execution strategy improves computational efficiency, while the default parameter configuration defines the main experimental setting used for fault simulation, known-drug evaluation, custom-target search, Pareto ranking, sensitivity analysis, and trajectory generation.

#### 4.9.1 Batched Vectorized Simulation

The number of required simulations grows combinatorially with the number of fault candidates, fault order, and intervention candidates. Therefore, simulations are executed using batched vectorized processing.

##### Algorithm 7

Pareto-based ranking and low-burden filtering

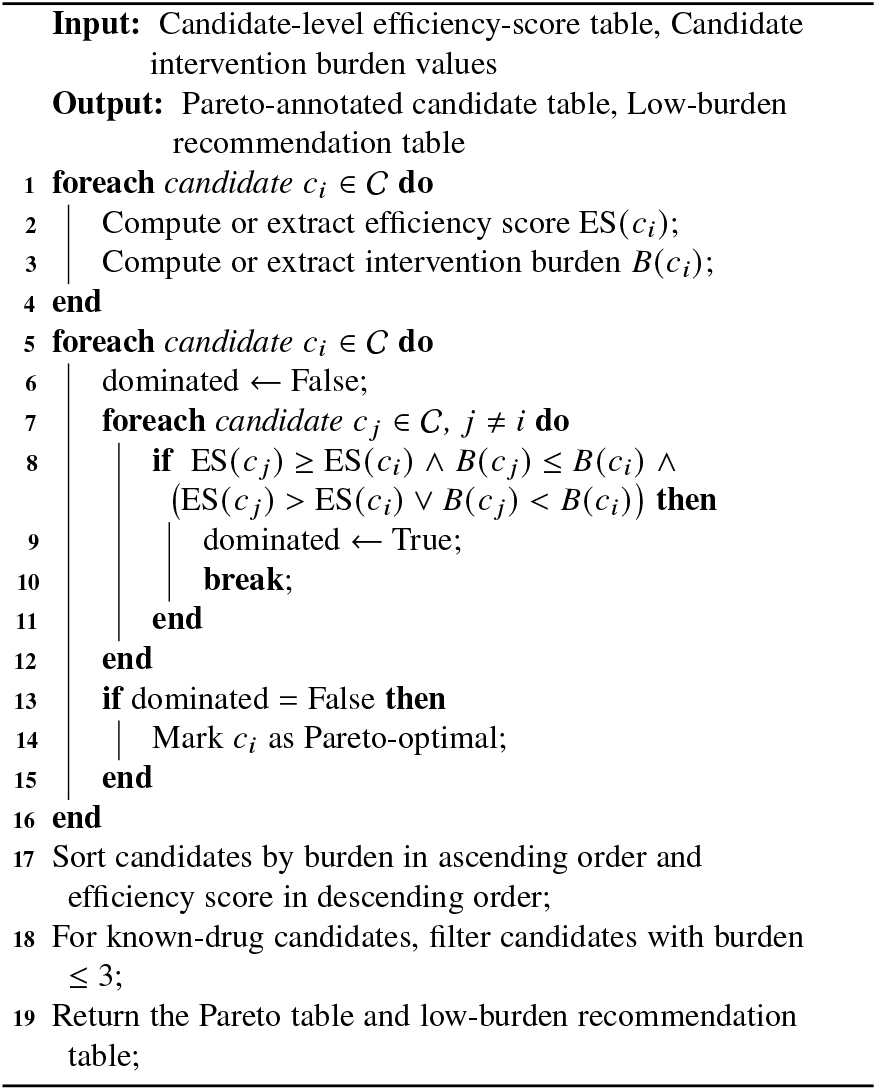

For a batch of *N*_*b*_ simulation conditions, node states are stored as arrays rather than scalar values. For each node *v*, the batch-level activation-probability vector at time *t* is represented as:

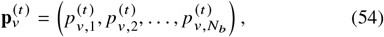

where 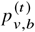 denotes the activation probability of node *v* at time *t* for the *b*-th simulation condition in the batch, with *b* = 1, 2, …, *N*_*b*_.

Each element in this array corresponds to one fault-only or fault-intervention simulation condition. Fault masks and intervention modes are also represented as arrays, allowing the same temporal DBN update equations to be applied simultaneously across all conditions in the batch. This avoids repeatedly evaluating the pathway one condition at a time and enables efficient large-scale simulation over fault combinations, drug vectors, and custom target pairs.

The implemented backend for the main analysis is vectorized NumPy. Therefore, runtime analyses are interpreted as comparisons between scalar or loop-based execution and batched vectorized execution. The default batch size is 1024 simulation conditions.

The complete number of simulation conditions generated by the main pipeline is summarized in Table 10.

**Table 10:**
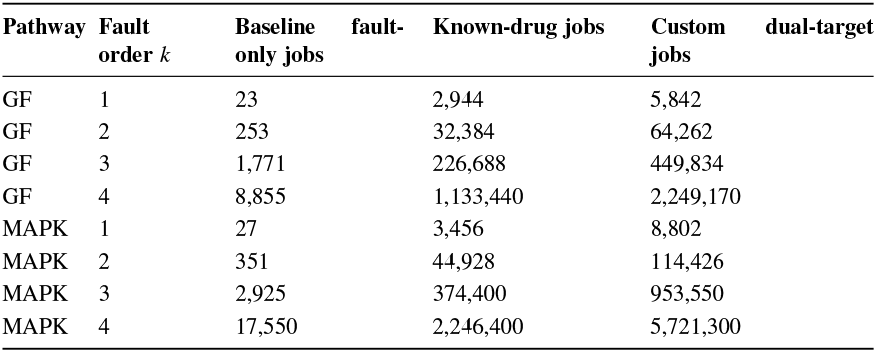
Number of simulation conditions evaluated by the main pipeline. Known-drug jobs include the no-drug baseline vector. Custom dual-target jobs include the empty custom-intervention baseline.

#### 4.9.2 Default Parameter Settings

The default parameters used in the main probabilistic DBN simulations are summarized in Table 11. These parameters define the main experimental configuration. Sensitivity analysis is performed later to evaluate whether candidate rankings remain stable under alternative parameter choices.

**Table 11:** Default probabilistic DBN parameter settings.

| Parameter | Default value | Meaning |
| --- | --- | --- |
| $\alpha$ | 0.95 | Maximum soft-logic propagation strength |
| $\eta$ | 0.05 | Basal activation leakage |
| $\lambda$ | 0.20 | Temporal persistence |
| $\rho$ | 0.95 | Fault penetrance |
| $\delta$ | 0.10 | Residual activity after inhibition or residual distance from activation |
| $T$ | 10 | Fixed temporal simulation horizon |
| $\epsilon$ | $10^{-4}$ | Output convergence tolerance |
| $x$ | 0.50 | Encoded-weight design parameter |
| Metformin mode | AMPK activation | Default Metformin intervention mode on $P_{20}$ |
| Backend | NumPy | Vectorized simulation backend |
| Batch size | 1024 | Number of simulation conditions processed per batch |

The parameters *α, η*, and *λ* control temporal DBN propagation. The parameters *ρ* and *δ* control fault penetrance and intervention strength, respectively. The parameter *T* defines the fixed simulation horizon, while *ϵ* is used to record convergence status. The encoded-weight parameter *x* balances transcription-factor/residual-protein imbalance and joint output-burden elevation. Backend and batch-size settings define the computational execution mode used in the main analysis.

## 5 Results

This section presents the findings obtained from probabilistic temporal DBN simulations of the GF and MAPK signaling pathways. The analyses encompass unique input vector selection, characterization of baseline multi-fault burden, evaluation of single-fault encoded-weight distributions, assessment of known-drug intervention strategies, Pareto-based ranking of intervention candidates, investigation of custom dual-target interventions, cross-pathway comparison of candidate performance, and computational runtime benchmarking.

### 5.1 Unique Input Vector Selection

An initial analysis was conducted to determine the input configuration that yields the lowest downstream output burden under fault-free, intervention-free probabilistic DBN conditions. Because both the GF and MAPK pathways contain five input nodes, an exhaustive evaluation of all 32 possible binary input vectors was performed.

For both pathways, the lowest-burden input vector was 00001. Under the default probabilistic DBN setting, this input vector produced a final total output burden of 2.6711 for GF and 3.5584 for MAPK. Therefore, the same input vector was used consistently for all subsequent baseline, multi-fault, known-drug, custom-target, Pareto, and runtime analyses.

Since the model is probabilistic, even in a fault-free situation, it produces nonzero output probabilities due to basal leakage, soft-logic propagation, and temporal persistence. Therefore, the selected input vector should not be interpreted as producing an all-zero output state. Instead, 00001 represents the lowest-burden input configuration under the selected DBN parameter setting.

### 5.2 Baseline Multi-fault Encoded Burden Before Intervention

Baseline simulations were performed for one, two, three, and four-fault scenarios without any intervention. Fig. 3 shows the resulting encoded-weight distributions for the GF and MAPK pathways.

**Fig. 3:**
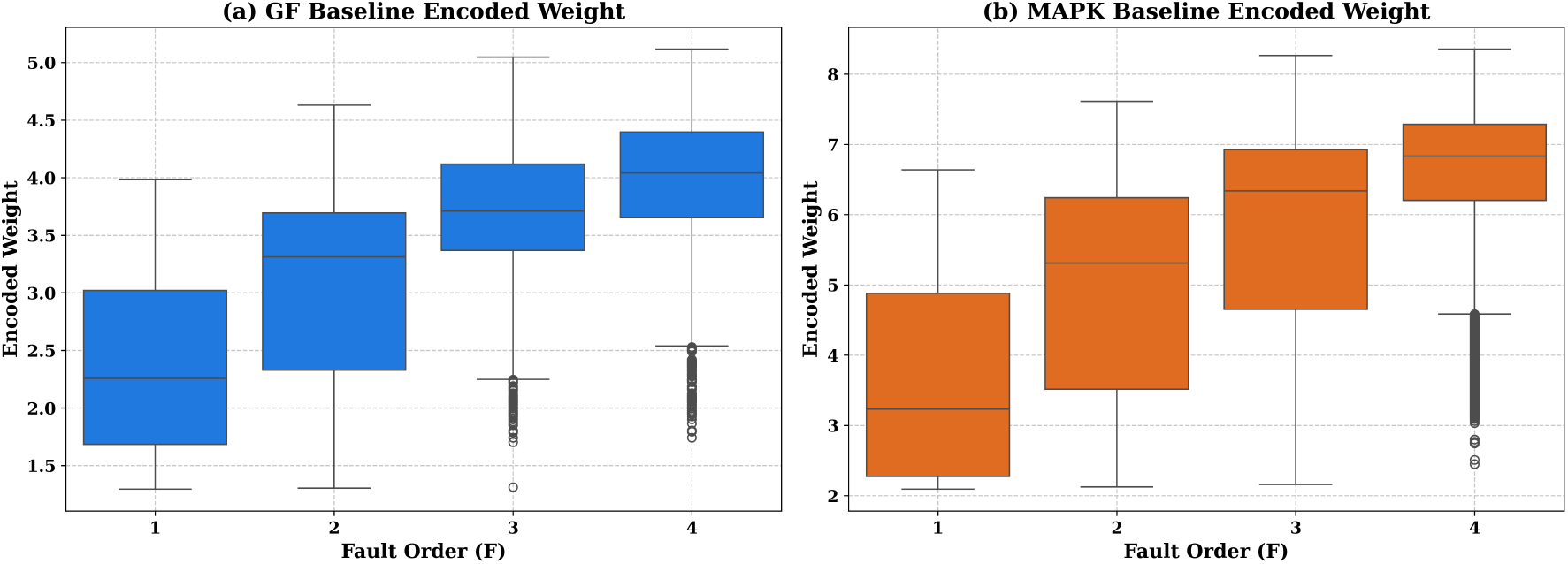
Baseline encoded-weight distributions under one, two, three, and four-fault scenarios for the GF and MAPK pathways

As illustrated in Fig. 3 and summarized quantitatively in Table 12, the encoded-weight distributions in both pathways shifted progressively upward as the number of simultaneous faults increased. For the GF pathway, the one-fault scenario exhibited the lowest baseline burden distribution, whereas the four-fault scenario produced the highest burden levels. A similar trend was observed for the MAPK pathway; however, the encoded-weight values were consistently higher than those of GF across all fault orders. These results indicate that the MAPK pathway generates a greater downstream burden than the GF pathway under comparable fault conditions.

**Table 12:** Baseline encoded-weight summary statistics across pathway-specific fault orders.

| Pathway | Fault order | No. of fault combinations | Minimum encoded weight | Q1 | Median | Q3 | Maximum encoded weight | Mean encoded weight |
| --- | --- | --- | --- | --- | --- | --- | --- | --- |
| GF | 1 | 23 | 1.2953 | 1.6837 | 2.2572 | 3.0197 | 3.9832 | 2.3525 |
| GF | 2 | 253 | 1.3032 | 2.3283 | 3.3117 | 3.6943 | 4.6300 | 3.1037 |
| GF | 3 | 1,771 | 1.3120 | 3.3688 | 3.7094 | 4.1170 | 5.0470 | 3.6061 |
| GF | 4 | 8,855 | 1.7424 | 3.6512 | 4.0414 | 4.3963 | 5.1156 | 3.9516 |
| MAPK | 1 | 27 | 2.0943 | 2.2739 | 3.2335 | 4.8806 | 6.6380 | 3.6425 |
| MAPK | 2 | 351 | 2.1253 | 3.5166 | 5.3114 | 6.2402 | 7.6124 | 4.9582 |
| MAPK | 3 | 2,925 | 2.1595 | 4.6535 | 6.3392 | 6.9252 | 8.2660 | 5.8758 |
| MAPK | 4 | 17,550 | 2.4505 | 6.2041 | 6.8329 | 7.2842 | 8.3547 | 6.5094 |

Fig. 3 further demonstrates that the severity of pathway disruption is not determined solely by the number of concurrent faults. Although higher-order fault scenarios generally resulted in larger encoded burdens, the distributions contained several lower-burden outliers, particularly in the three- and four-fault cases. The corresponding summary statistics in Table 12 support this observation by showing substantial variability within each fault order. This variability suggests that the encoded burden depends not only on the fault order itself but also on the specific locations and combinations of faulty nodes within the pathway topology.

### 5.3 Single-Fault Intervention Response Profiles

Single-fault encoded-weight plots were generated to evaluate the effects of the known-drug vectors across individual fault scenarios. Fig. 4 presents the GF pathway results, while Fig. 5 presents the corresponding MAPK pathway results.

**Fig. 4:**
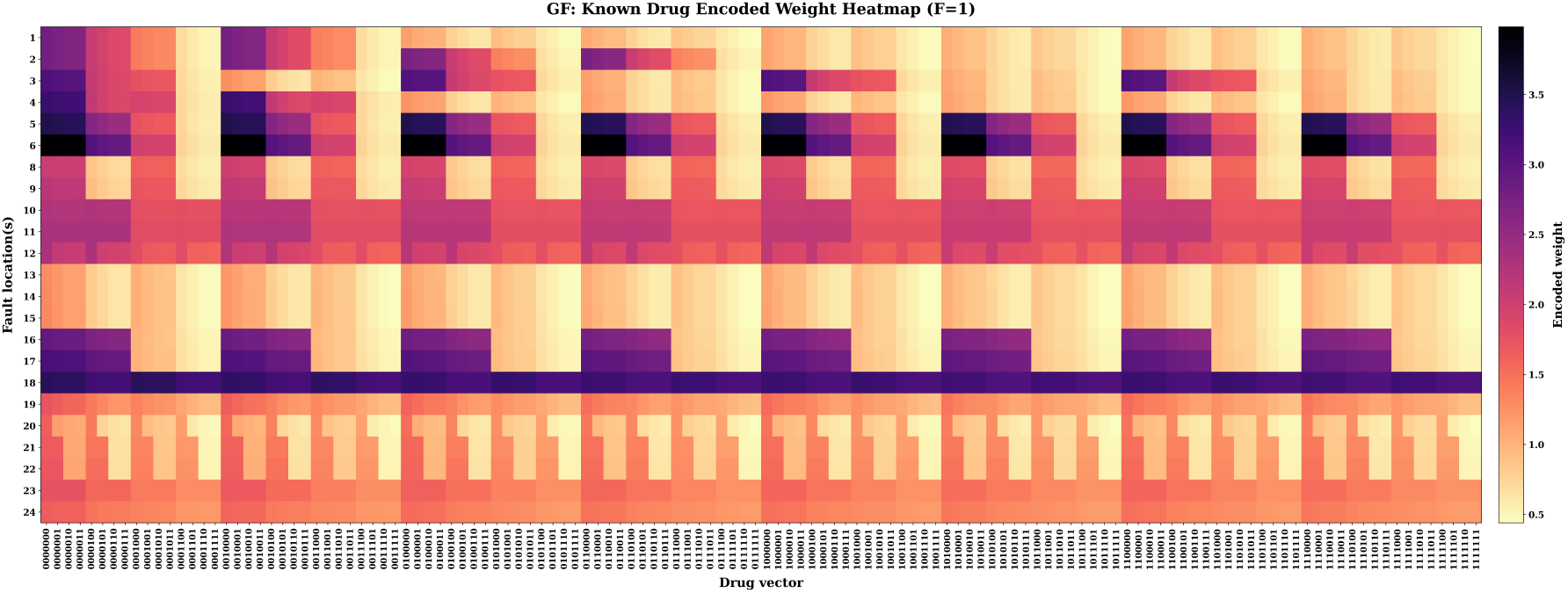
Encoded-weight heatmap for GF single-fault scenarios evaluated across all 128 known-drug intervention vectors. Each row corresponds to a distinct single-fault condition in the GF pathway, and each column represents one known-drug vector. Cell colours indicate the resulting encoded-weight burden after intervention, with lighter colours (e.g., yellow) representing lower encoded-weight values and therefore more effective burden reduction, and darker colours (e.g., black) representing higher encoded-weight values and less effective intervention performance. Structured colour patterns reveal how the effectiveness of specific drug combinations varies across different fault locations, highlighting intervention vectors that consistently maintain low encoded burden across multiple single-fault scenarios.

**Fig. 5:**
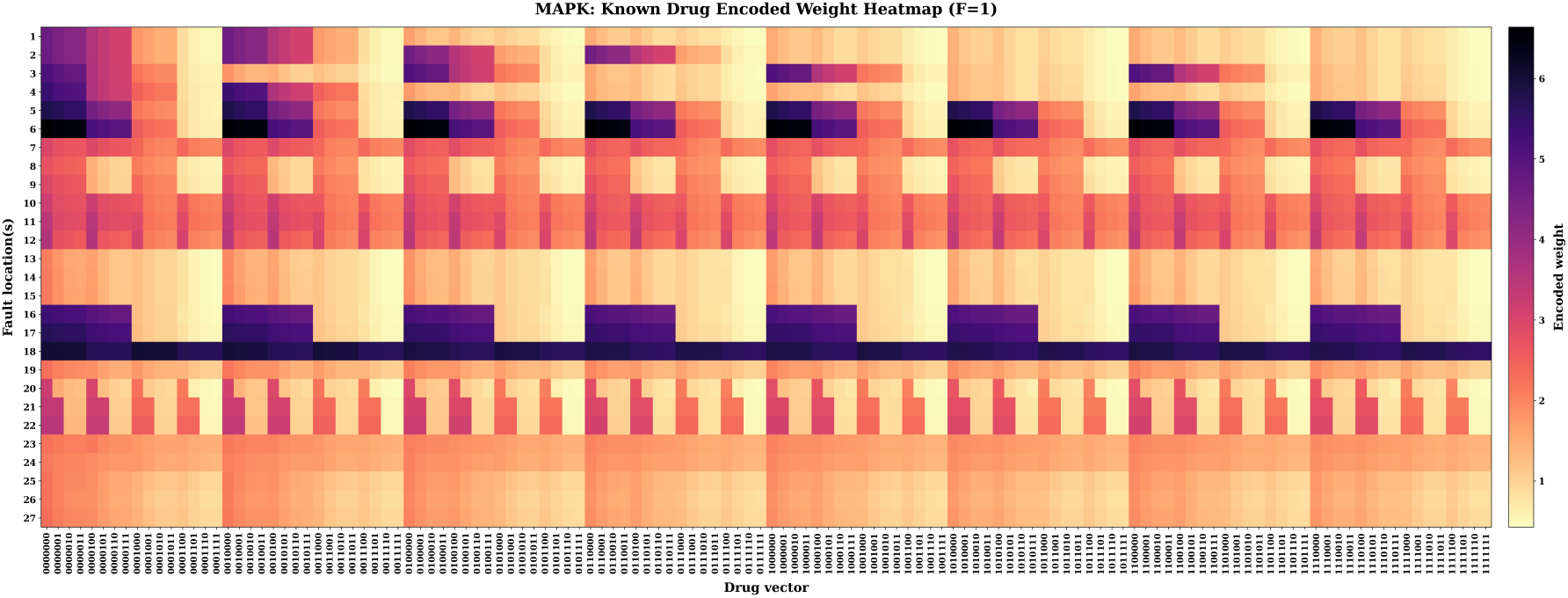
Encoded-weight heatmap for MAPK single-fault scenarios evaluated across all 128 known-drug intervention vectors. Each row corresponds to a distinct single-fault condition in the MAPK pathway, and each column represents one known-drug vector. Cell colours indicate the resulting encoded-weight burden after intervention, with lighter colours (e.g., yellow) representing lower encoded-weight values and therefore more effective burden reduction, and darker colours (e.g., black) representing higher encoded-weight values and less effective intervention performance. Structured colour patterns reveal how the effectiveness of specific drug combinations varies across different fault locations, highlighting intervention vectors that consistently maintain low encoded burden across multiple single-fault scenarios.

Both heatmaps showed structured bands of low and high encoded burden across the drug-vector space. This indicates that drug-vector performance was not random, but strongly dependent on which pathway targets were included in the combination.

Several drug-vector groups consistently reduced encoded weight across many single-fault scenarios, while other vectors produced higher residual burden.

As evident from Fig. 5, the MAPK single-fault heatmap showed broader high-burden regions than the GF heatmap, which is consistent with the higher baseline burden observed in Fig. 3. These results also show why aggregate efficiency scoring is necessary: a drug vector may perform well for several fault locations but not for every single-fault condition. Therefore, intervention candidates were ranked using their burden reduction across all fault combinations of a given fault order, rather than by visual inspection of one fault-response pattern.

### 5.4 Known-drug Efficiency Across Fault Orders

The efficiency of all 128 known-drug vectors was evaluated under one, two, three, and four-fault conditions in both the GF and MAPK pathways. Figs. 6-9 shows the resulting efficiency-score distributions for each fault order.

**Fig. 6:**
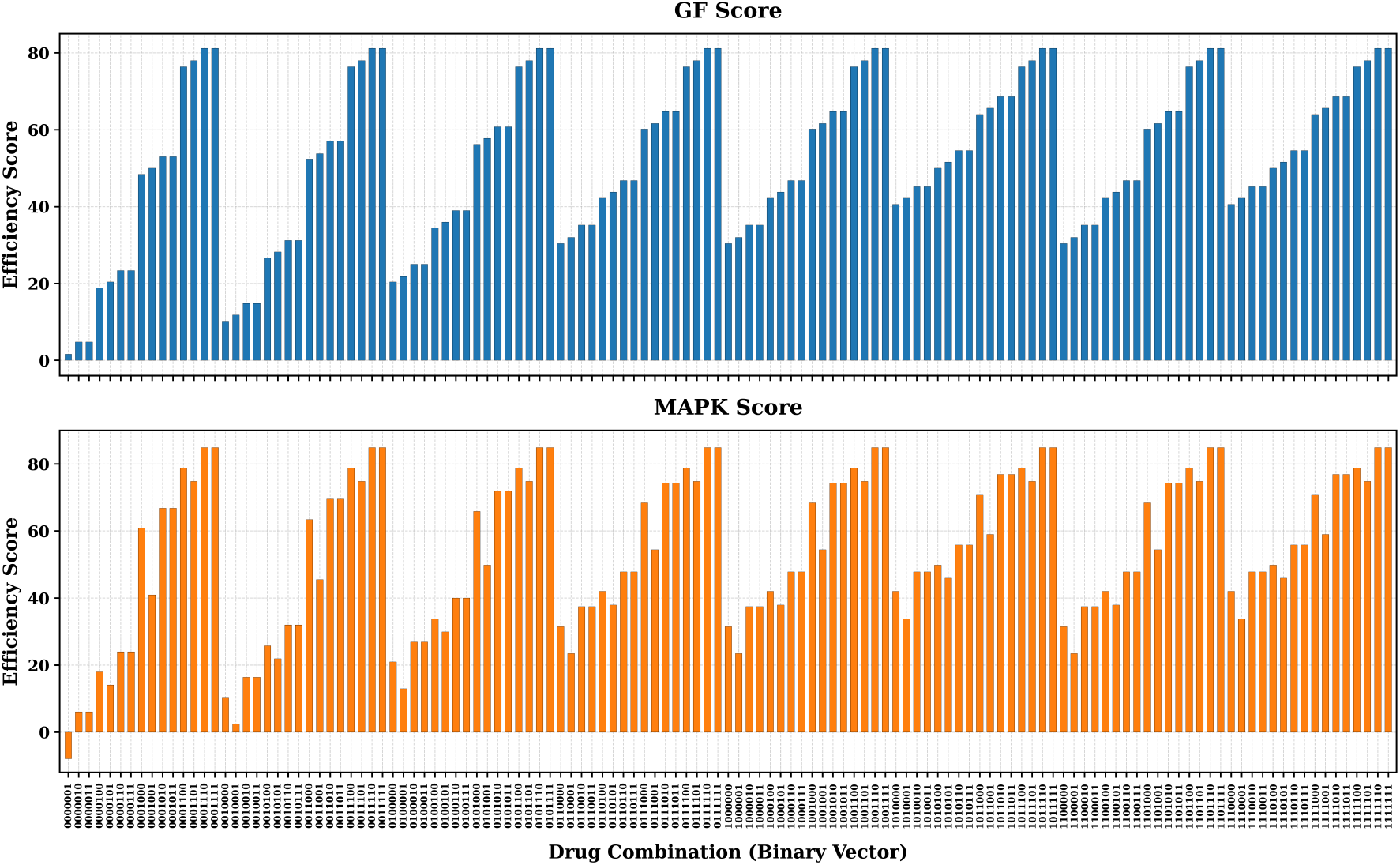
Known-drug efficiency scores for the GF and MAPK pathways under one-fault scenarios. Each point represents the efficiency score achieved by a specific known-drug intervention vector when evaluated against all single-fault conditions in the probabilistic temporal DBN framework. The figure compares the relative performance of the 128 candidate drug vectors, highlighting differences in burden reduction capability between the two pathways and identifying high-performing intervention combinations that consistently achieve strong fault-mitigation effects.

**Fig. 7:**
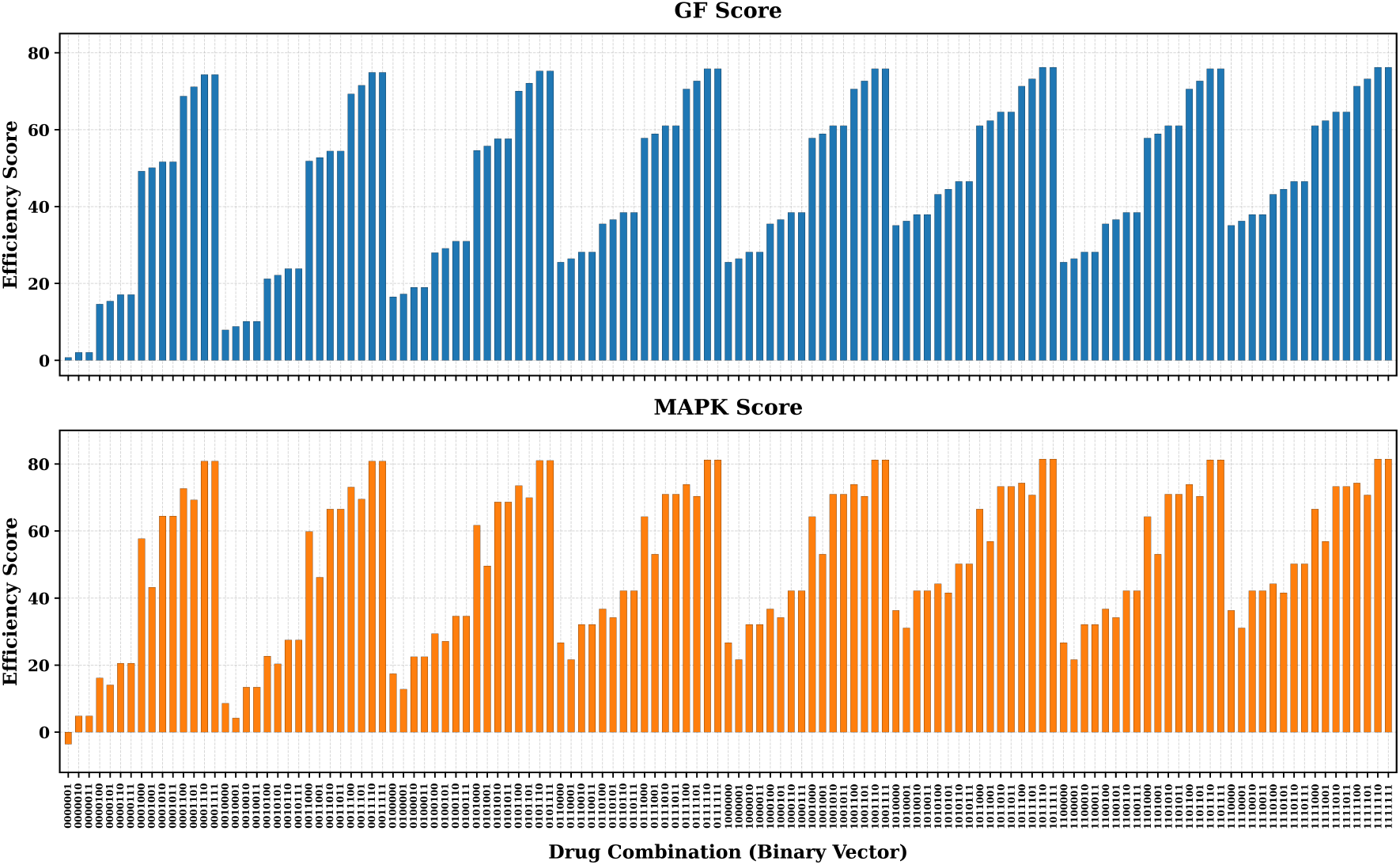
Known-drug efficiency scores for the GF and MAPK pathways under two-fault scenarios. Each point represents the efficiency score achieved by a specific known-drug intervention vector when evaluated against all two-fault conditions in the probabilistic temporal DBN framework. The figure compares the relative performance of the 128 candidate drug vectors, highlighting differences in burden reduction capability between the two pathways and identifying high-performing intervention combinations that consistently achieve strong fault-mitigation effects.

**Fig. 8:**
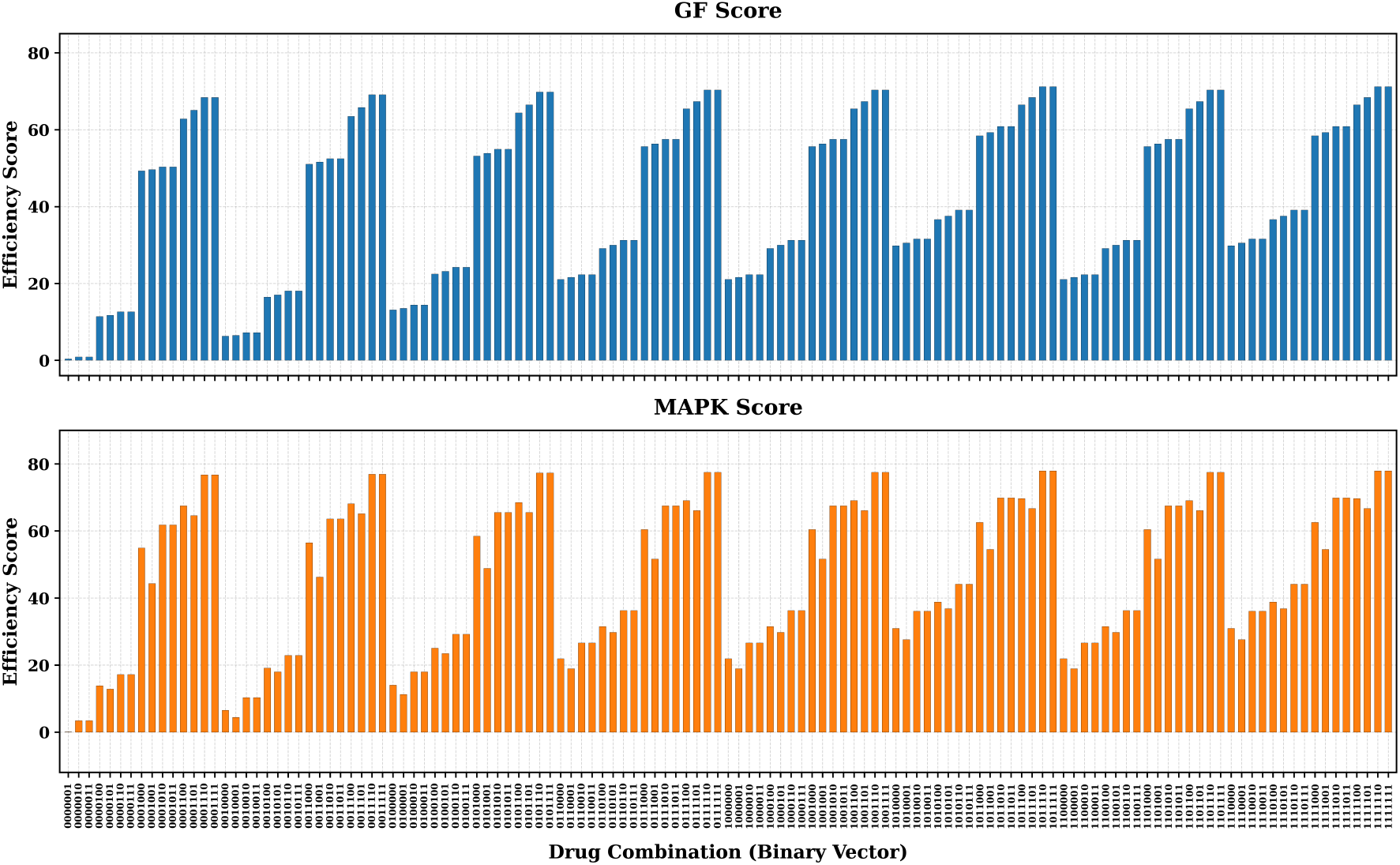
Known-drug efficiency scores for the GF and MAPK pathways under three-fault scenarios. Each point represents the efficiency score achieved by a specific known-drug intervention vector when evaluated against all three-fault conditions in the probabilistic temporal DBN framework. The figure compares the relative performance of the 128 candidate drug vectors, highlighting differences in burden reduction capability between the two pathways and identifying high-performing intervention combinations that consistently achieve strong fault-mitigation effects.

**Fig. 9:**
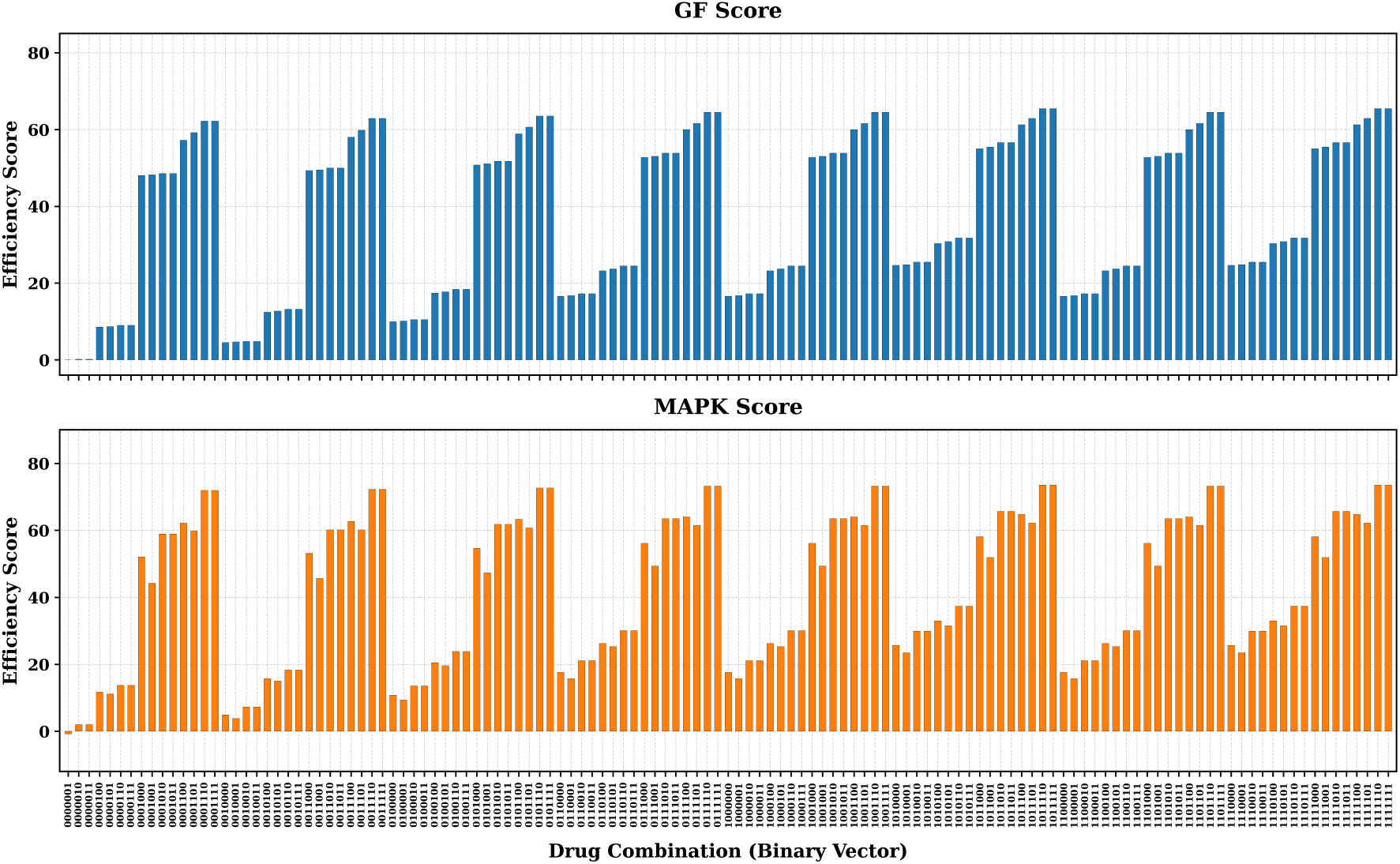
Known-drug efficiency scores for the GF and MAPK pathways under four-fault scenarios. Each point represents the efficiency score achieved by a specific known-drug intervention vector when evaluated against all four-fault conditions in the probabilistic temporal DBN framework. The figure compares the relative performance of the 128 candidate drug vectors, highlighting differences in burden reduction capability between the two pathways and identifying high-performing intervention combinations that consistently achieve strong fault-mitigation effects.

Across all four fault orders, the efficiency-score plots showed repeated structured patterns. Drug vectors containing downstream pathway inhibitors generally produced higher efficiency scores, whereas several low-burden or upstream-only combinations produced weaker burden reduction.

For every fault order in both pathways, the best low-burden known-drug vector was U0126+LY294002+Temsirolimus. This combination has an intervention burden of 3. The maximum-efficiency vector was Lapatinib+AG1024+U0126+LY294002+Temsirolimus+Metformin, with an intervention burden of 6. The efficiency scores and burden values for these leading intervention candidates across all fault orders are summarized in Table 13.

**Table 13:**
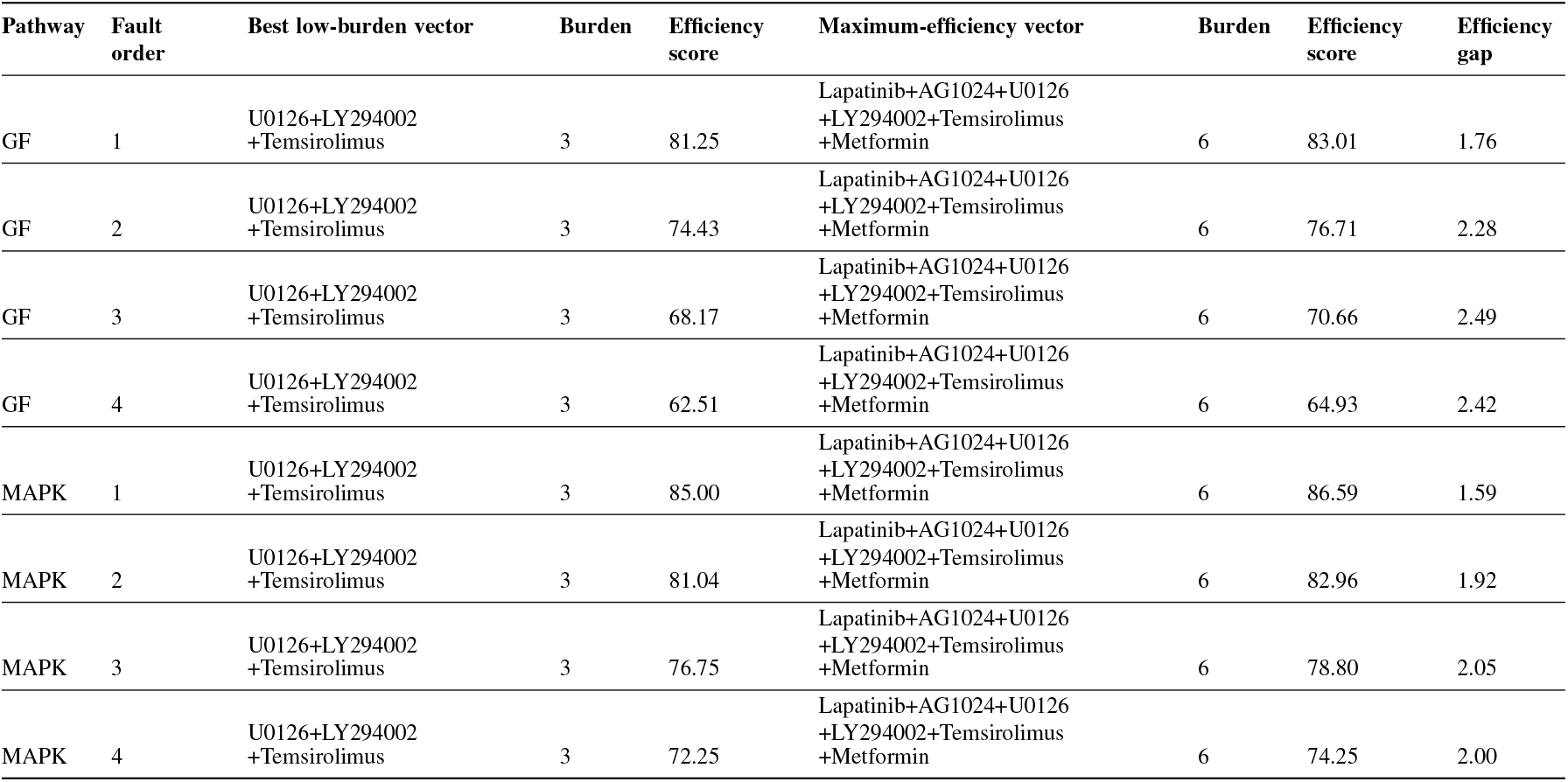
Best low-burden and maximum-efficiency known-drug vectors across pathways and fault orders.

As shown in Table 13, the three-drug vector U0126+LY294002+Temsirolimus remained close to the six-drug maximum-efficiency vector across all fault orders. In GF, the efficiency gap ranged from 1.76 percentage points in the one-fault scenario to 2.49 percentage points in the three-fault scenario, while in MAPK, the gap ranged from 1.59 to 2.05 percentage points. Therefore, the three-drug low-burden vector achieved most of the burden reduction produced by the maximum-efficiency vector while requiring only half the number of drugs.

Table 13 also shows that the efficiency score decreased as the number of concurrent faults increased. For the best low-burden vector in GF, the efficiency score declined from 81.25 in the one-fault scenario to 62.51 in the four-fault scenario. In MAPK, the corresponding score decreased from 85.00 to 72.25. A similar trend was observed for the maximum-efficiency vector, whose score decreased from 83.01 to 64.93 in GF and from 86.59 to 74.25 in MAPK. These results indicate that increasing multi-fault dysregulation reduces the relative effectiveness of known-drug intervention.

### 4.5 Pathway-specific Known-drug Efficiency Profiles

To further examine how intervention performance changed with increasing fault order, known-drug efficiency scores were plotted separately for the GF and MAPK pathways across the one, two, three, and four-fault scenarios. The GF results are shown in Fig. 10, while the corresponding MAPK results are shown in Fig. 11.

**Fig. 10:**
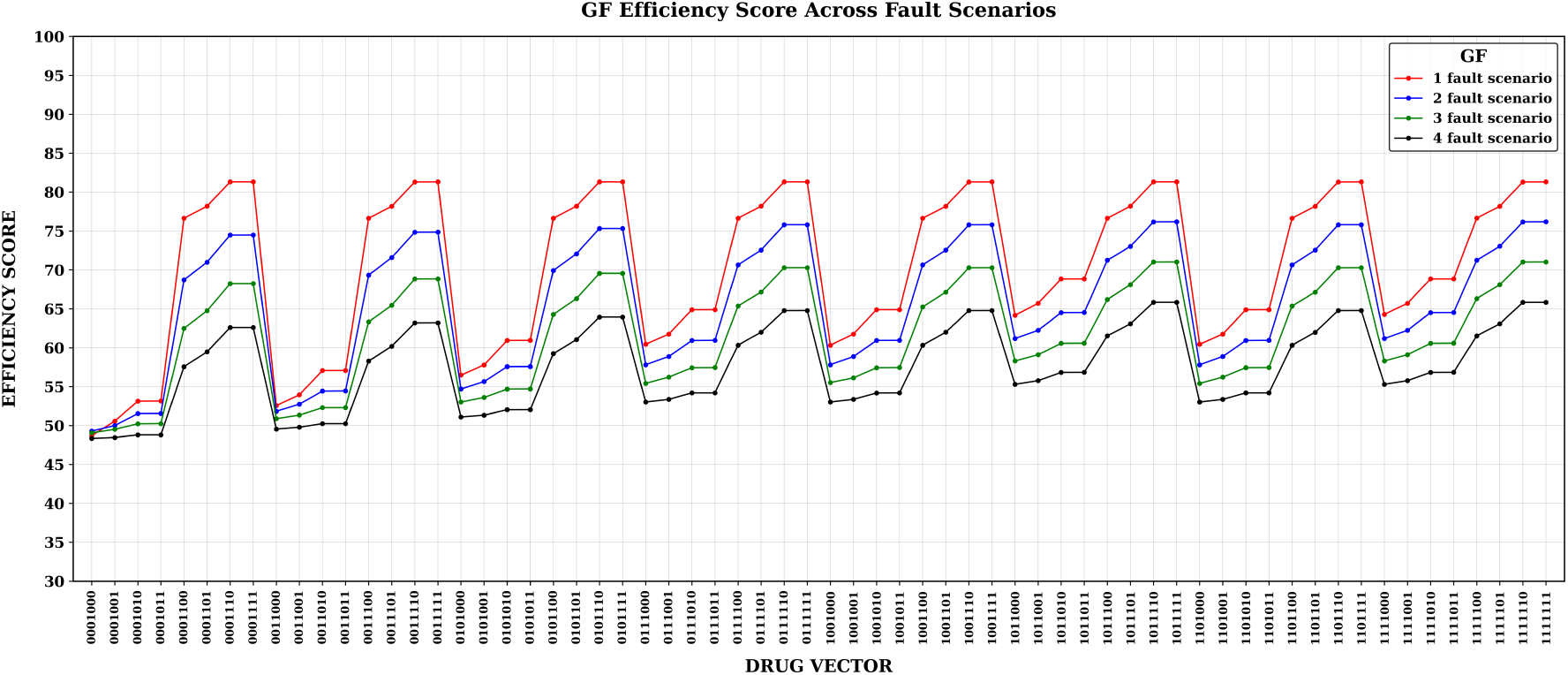
GF known-drug efficiency scores across one, two, three, and four-fault scenarios.

**Fig. 11:**
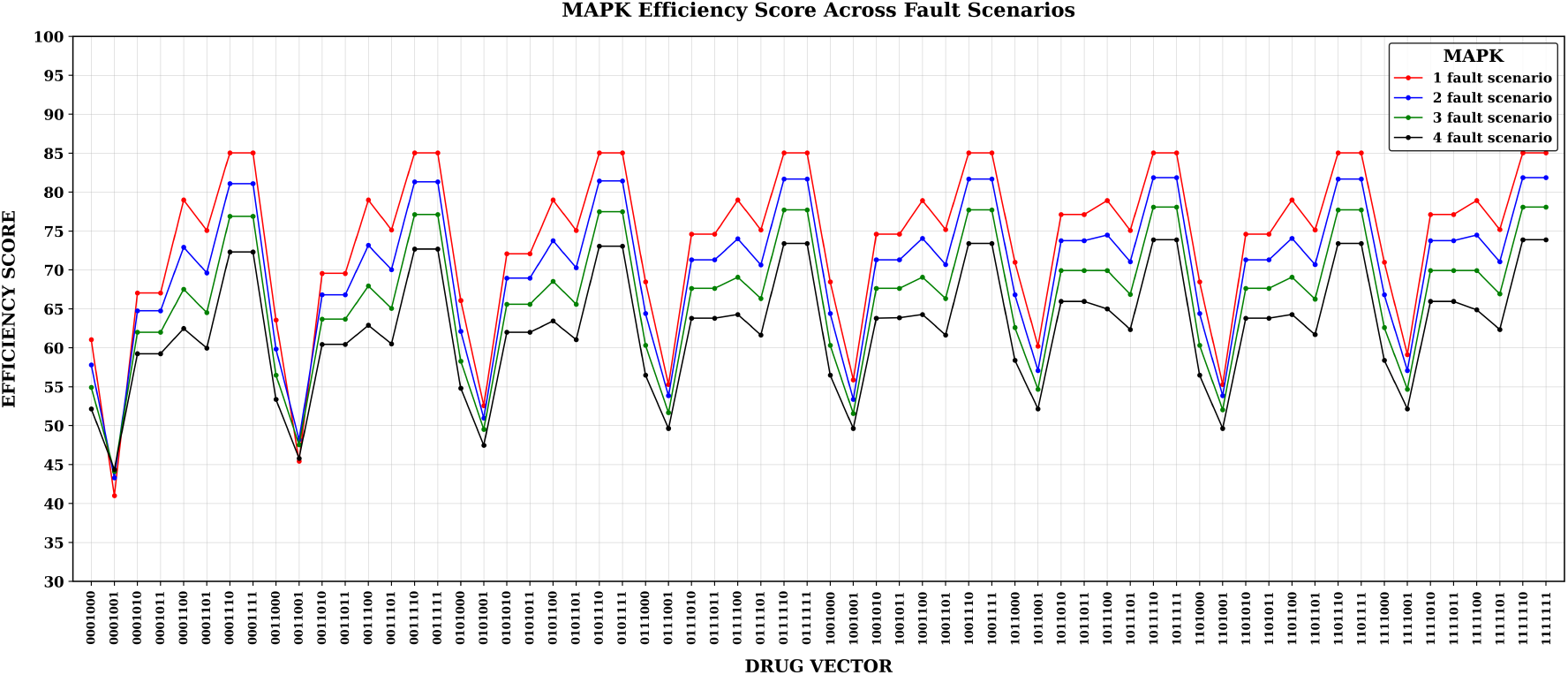
MAPK known-drug efficiency scores across one, two, three, and four-fault scenarios.

Both pathway-specific plots, shown in Figs. 10 and 11 demonstrated a consistent reduction in efficiency score as the fault order increased. Although the absolute efficiency values were higher in MAPK than in GF, the same broad ranking structure was preserved across the four fault orders in both figures. This suggests that the strongest known drug candidates remained stable even as the number of simultaneous faults increased.

The decrease in efficiency was broadly comparable between pathways. As shown in Table 14, the best low-burden vector exhibited an absolute efficiency reduction of 13.42 points in GF and 12.90 points in MAPK when moving from one-fault to four-fault scenarios, while the maximum-efficiency vector decreased by 12.76 and 12.49 points, respectively. These results demonstrate a consistent decline in intervention effectiveness as the number of concurrent faults increases, suggesting that the reduction in known-drug performance with increasing fault order is a general feature of the multi-fault simulation setting rather than a pathway-specific effect.

**Table 14:** Efficiency decrease from one-fault to four-fault scenarios.

| Pathway | Candidate type | $F = 1$ | $F = 2$ | $F = 3$ | $F = 4$ | Absolute decrease from $F = 1$ to $F = 4$ |
| --- | --- | --- | --- | --- | --- | --- |
| GF | Best low-burden vector | 81.25 | 74.43 | 68.17 | 62.51 | 18.74 |
| GF | Maximum-efficiency vector | 83.01 | 76.71 | 70.66 | 64.93 | 18.08 |
| MAPK | Best low-burden vector | 85.00 | 81.04 | 76.75 | 72.25 | 12.75 |
| MAPK | Maximum-efficiency vector | 86.59 | 82.96 | 78.80 | 74.25 | 12.34 |

### 4.6 Pareto Ranking of Known-drug Candidates

Pareto analysis was performed to evaluate the trade-off between intervention efficiency and intervention burden. The two optimization objectives were defined as maximizing the intervention efficiency score while simultaneously minimizing intervention burden, measured as the number of drugs included in the intervention vector. Each known-drug vector was therefore represented by these two competing objective values. Fig. 12 presents the Pareto-front analysis for the GF and MAPK pathways across one, two, three, and four-fault scenarios.

**Fig. 12:**
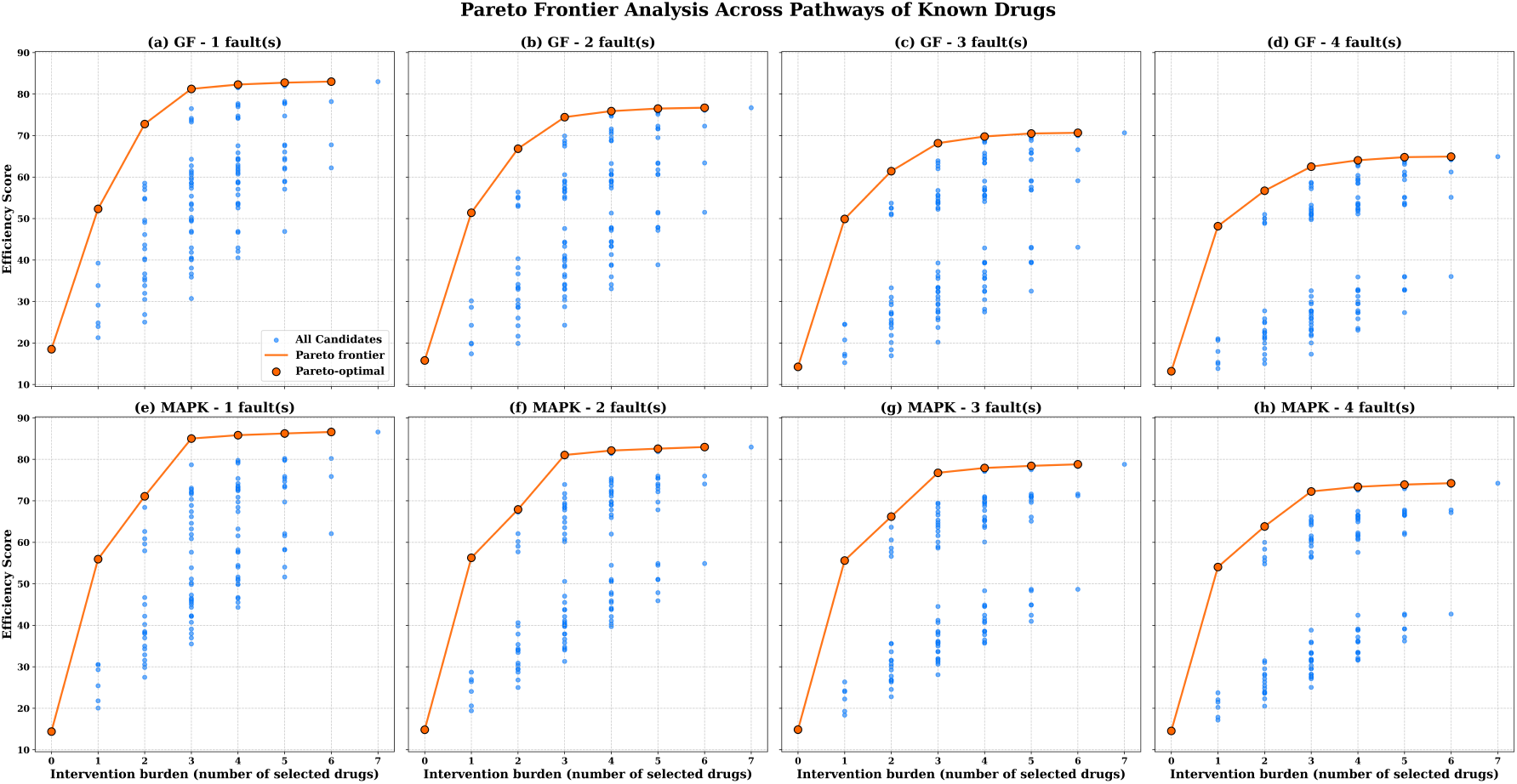
Pareto-front analysis of known-drug candidates across the GF and MAPK pathways under one, two, three, and four-fault scenarios. The eight panels show efficiency score versus intervention burden for all known-drug vectors. Pareto-optimal candidates form the upper frontier of each panel, representing the best trade-offs between burden and efficiency. The plots highlight low-burden high-efficiency candidates, including U0126+LY294002+Temsirolimus, and show that efficiency gains become smaller as additional drugs are added.

The Pareto-front plots showed that efficiency increased sharply from burden 0 to burden 3, after which additional drugs produced smaller gains. This pattern was visible in both pathways and across all fault orders. The three-drug vector U0126+LY294002+Temsirolimus was consistently positioned near the upper region of the Pareto front, while the six-drug vector achieved the maximum efficiency score.

As summarized in Table 15, Pareto-optimal candidates were identified across multiple intervention burden levels for both pathways and all fault orders. The table provides a numerical summary of the candidates defining the Pareto front and complements the visual trends observed in Fig. 12. The three-drug vector U0126+LY294002+Temsirolimus repeatedly appeared among the strongest low-burden Pareto-optimal solutions, while the six-drug combination remained the maximum-efficiency candidate.

**Table 15:**
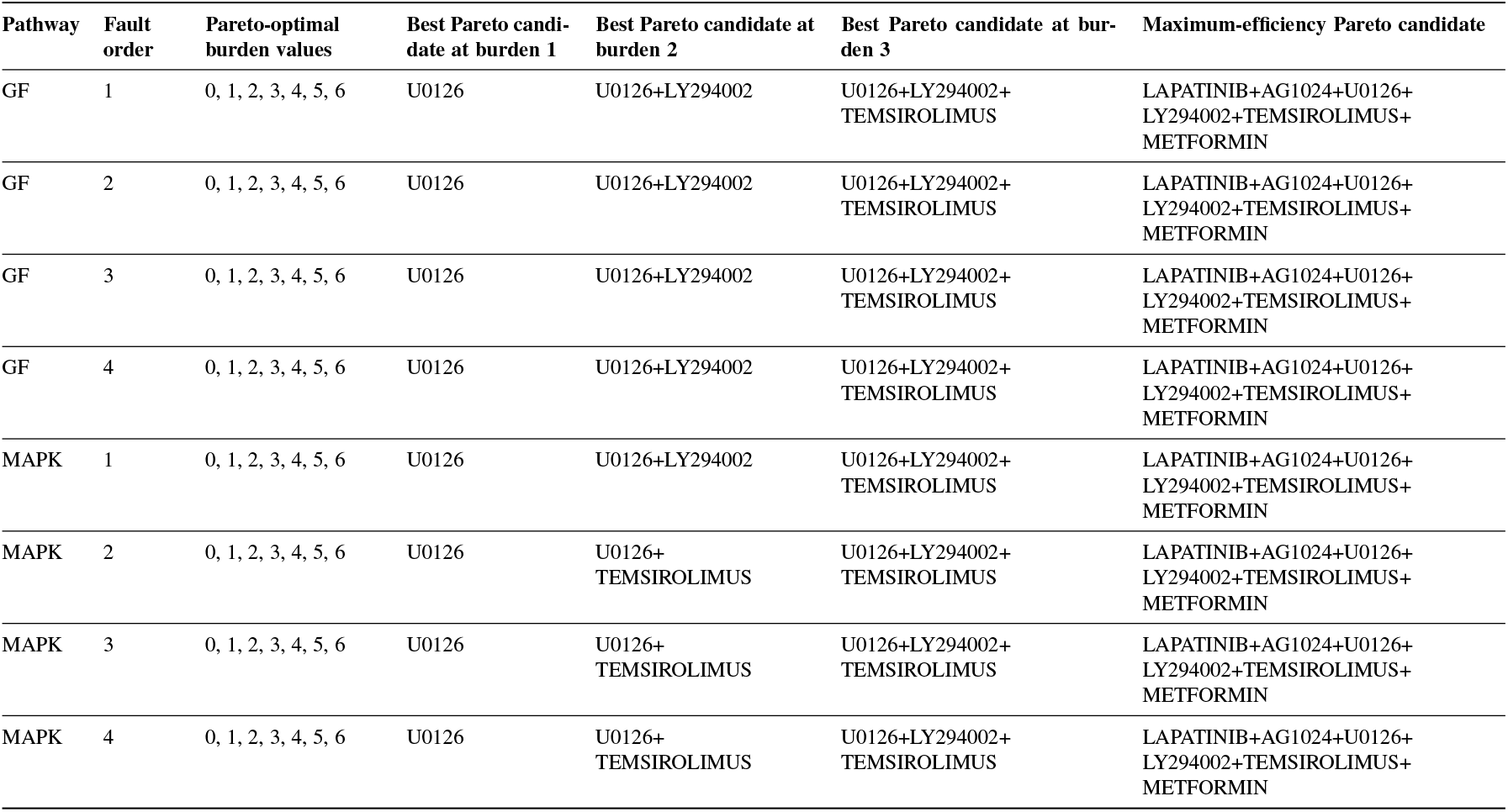
Pareto-optimal known-drug candidates across pathways and fault orders.

| Pathway | Fault order | Pareto-optimal burden values | Best Pareto candidate at burden 1 | Best Pareto candidate at burden 2 | Best Pareto candidate at burden 3 | Maximum-efficiency Pareto candidate |
| --- | --- | --- | --- | --- | --- | --- |
| GF | 1 | 0, 1, 2, 3, 4, 5, 6 | U0126 | U0126+LY294002 | U0126+LY294002+TEMSIROLIMUS | LAPATINIB+AG1024+U0126+LY294002+TEMSIROLIMUS+METFORMIN |
| GF | 2 | 0, 1, 2, 3, 4, 5, 6 | U0126 | U0126+LY294002 | U0126+LY294002+TEMSIROLIMUS | LAPATINIB+AG1024+U0126+LY294002+TEMSIROLIMUS+METFORMIN |
| GF | 3 | 0, 1, 2, 3, 4, 5, 6 | U0126 | U0126+LY294002 | U0126+LY294002+TEMSIROLIMUS | LAPATINIB+AG1024+U0126+LY294002+TEMSIROLIMUS+METFORMIN |
| GF | 4 | 0, 1, 2, 3, 4, 5, 6 | U0126 | U0126+LY294002 | U0126+LY294002+TEMSIROLIMUS | LAPATINIB+AG1024+U0126+LY294002+TEMSIROLIMUS+METFORMIN |
| MAPK | 1 | 0, 1, 2, 3, 4, 5, 6 | U0126 | U0126+LY294002 | U0126+LY294002+TEMSIROLIMUS | LAPATINIB+AG1024+U0126+LY294002+TEMSIROLIMUS+METFORMIN |
| MAPK | 2 | 0, 1, 2, 3, 4, 5, 6 | U0126 | U0126+TEMSIROLIMUS | U0126+LY294002+TEMSIROLIMUS | LAPATINIB+AG1024+U0126+LY294002+TEMSIROLIMUS+METFORMIN |
| MAPK | 3 | 0, 1, 2, 3, 4, 5, 6 | U0126 | U0126+TEMSIROLIMUS | U0126+LY294002+TEMSIROLIMUS | LAPATINIB+AG1024+U0126+LY294002+TEMSIROLIMUS+METFORMIN |
| MAPK | 4 | 0, 1, 2, 3, 4, 5, 6 | U0126 | U0126+TEMSIROLIMUS | U0126+LY294002+TEMSIROLIMUS | LAPATINIB+AG1024+U0126+LY294002+TEMSIROLIMUS+METFORMIN |

Fig. 12 shows that efficiency generally increased with intervention burden, although the rate of improvement varied across pathways and fault orders. The Pareto-front plots provide a compact summary of the efficiency–burden relationship for the evaluated known-drug candidates.

### 4.7 Known-drug and Custom-target Candidate Comparison

The known-versus-custom comparison was performed to compare high-performing known-drug vectors with top-ranked custom dual-target candidates. The GF comparison is shown in Fig. 13, and the MAPK comparison is shown in Fig. 14.

**Fig. 13:**
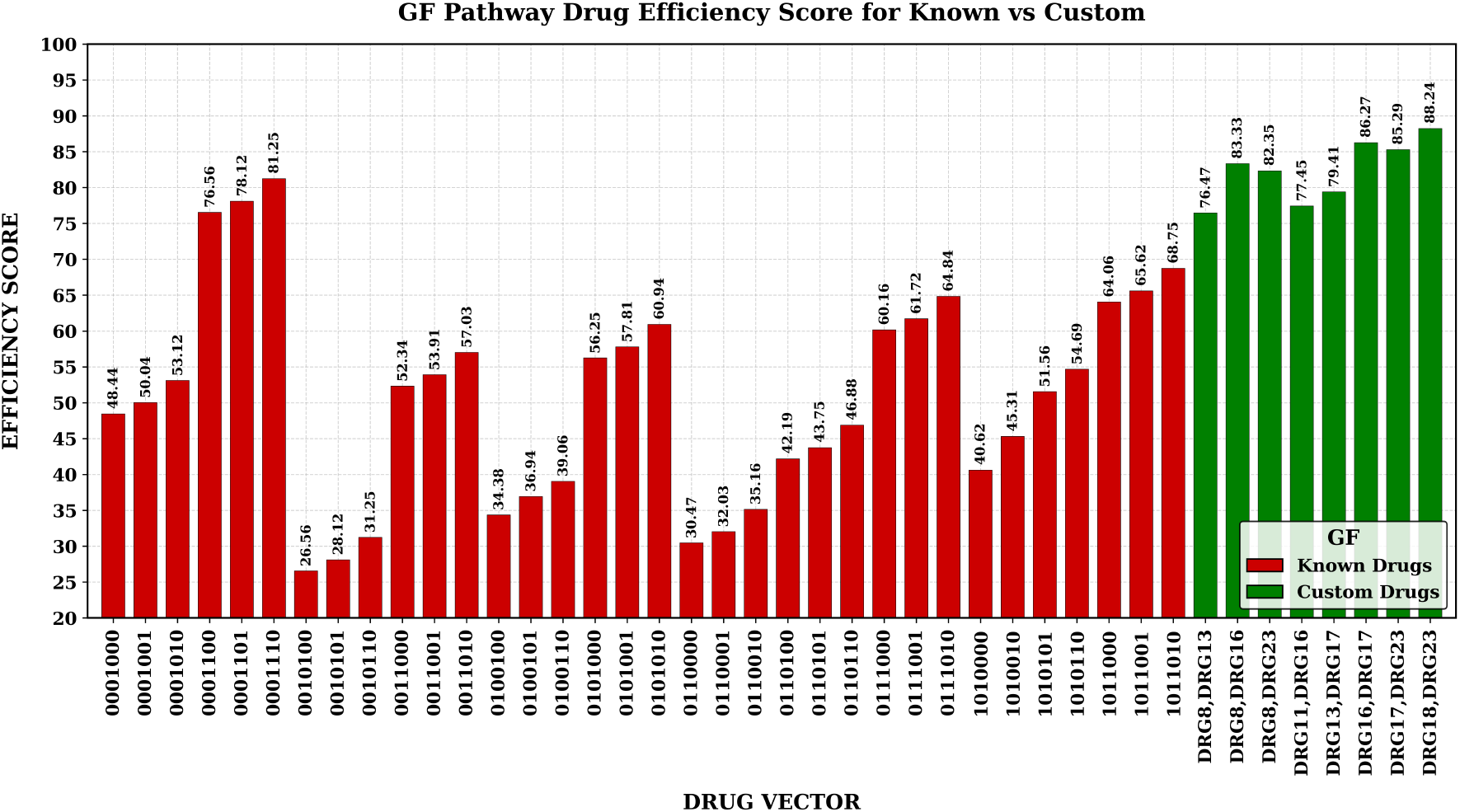
Efficiency-score comparison between selected known-drug vectors and custom dual-target candidates in the GF pathway

**Fig. 14:**
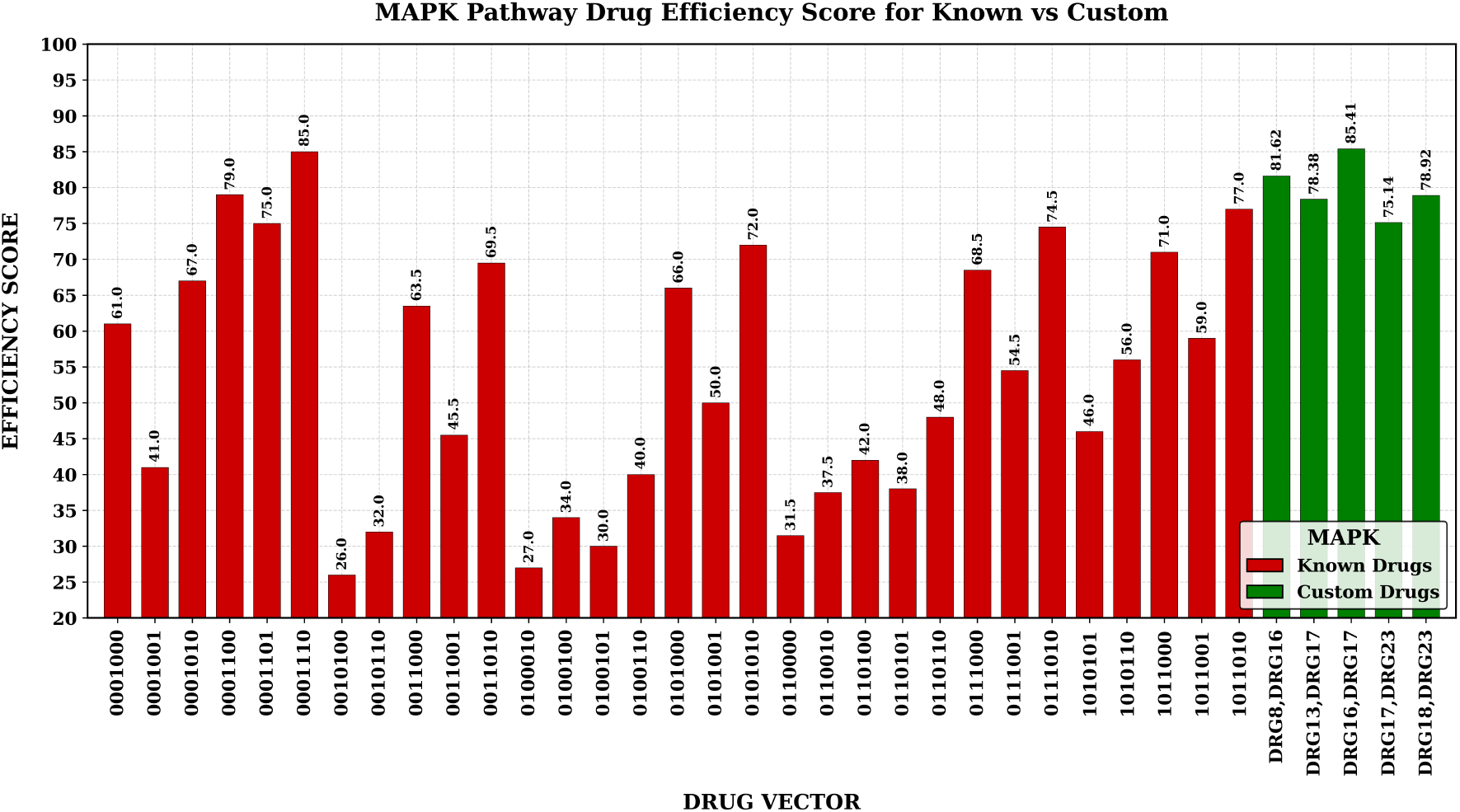
Efficiency-score comparison between selected known-drug vectors and custom dual-target candidates in the MAPK pathway

In the GF pathway, as shown in Fig. 13, the highest custom-target score was obtained by [DRG18,DRG23], with an efficiency score of 88.24. Here, DRG18 and DRG23 refer to internal pathway target identifiers used in the custom intervention search; therefore, [DRG18,DRG23] represent the simultaneous inhibition of those two pathway nodes rather than a specific approved drug combination. The same figure shows that the next highest custom candidates were [DRG16,DRG17] (86.27), [DRG17,DRG23] (85.29), and [DRG8,DRG16] (83.33). Among the known-drug vectors included in the comparison, Fig. 13 indicates that 0001110 achieved the highest efficiency score, reaching 81.25.

In the MAPK pathway, as evident from Fig. 14, the highest custom-target score was obtained by [DRG16,DRG17], with an efficiency score of 85.41. The next highest custom candidates were [DRG8,DRG16] with 81.62, [DRG18,DRG23] with 78.92, [DRG13,DRG17] with 78.38, and [DRG17,DRG23] with 75.14. The strongest known-drug vector displayed in this comparison was 0001110, with an efficiency score of 85.00.

These comparisons show that custom dual-target candidates can reach or exceed the efficiency of the best selected known-drug vectors. As summarized in Table 16, the highest-scoring custom candidate in GF achieved an efficiency score of 88.24 compared with 81.25 for the best known-drug vector shown in the comparison, corresponding to an improvement of 6.99 percentage points. In MAPK, the highest-scoring custom candidate achieved an efficiency score of 85.41 compared with 85.00 for the best known-drug vector shown in the comparison, corresponding to an improvement of 0.41 percentage points.

**Table 16:**
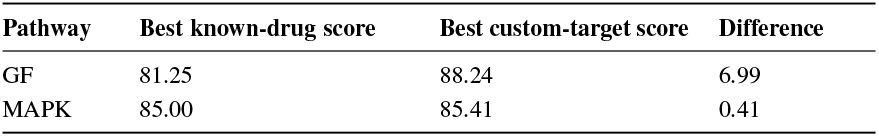
Best known-drug versus best custom-target candidate in the known/custom comparison plots.

These results indicate that the custom dual-target search is particularly useful for identifying high-impact pathway intervention points. However, these custom candidates should be interpreted as computational inhibition-point hypotheses, not as validated drug combinations.

### 5.8 Cross-pathway Candidate Comparison

To compare candidate behaviour across pathways, the high-efficiency known-drug and custom-target candidates were summarized in a combined GF/MAPK comparison plot, shown in Fig. 15.

**Fig. 15:**
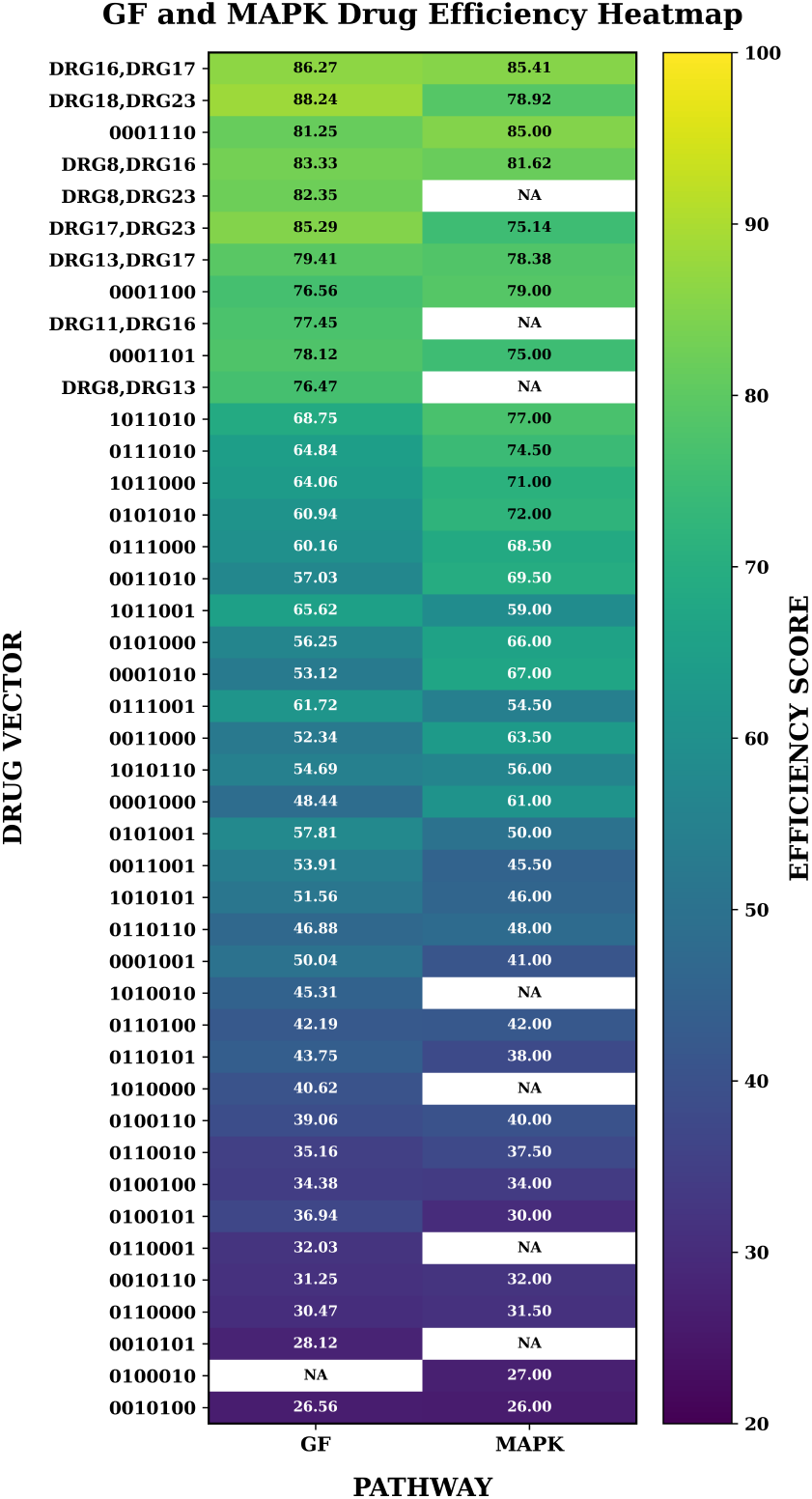
Cross-pathway efficiency heatmap comparing selected known-drug vectors and custom dual-target candidates in GF and MAPK

Fig. 15 shows that several candidates achieved high efficiency in both pathways, while others displayed more pathway-specific performance. Based on the values shown in the figure, the custom target pair [DRG16,DRG17] performed strongly in both GF and MAPK, with efficiency scores of 86.27 and 85.41, respectively. [DRG18,DRG23] produced the highest GF score, 88.24, but a lower MAPK score of 78.92. The known-drug vector 0001110 also performed strongly in both pathways, with a GF score of 81.25 and a MAPK score of 85.00.

The values presented in Fig. 15 indicate two main observations. First, several custom target pairs achieved efficiency scores comparable to or higher than those of the known-drug vectors in one or both pathways. Second, candidate performance was not identical across GF and MAPK, suggesting that pathway topology and downstream output structure influence intervention effectiveness.

### 5.9 Computational Performance

Computational performance was evaluated by comparing scalar/simple-loop execution with batched vectorized NumPy execution. The runtime comparison is shown in Fig. 16.

**Fig. 16:**
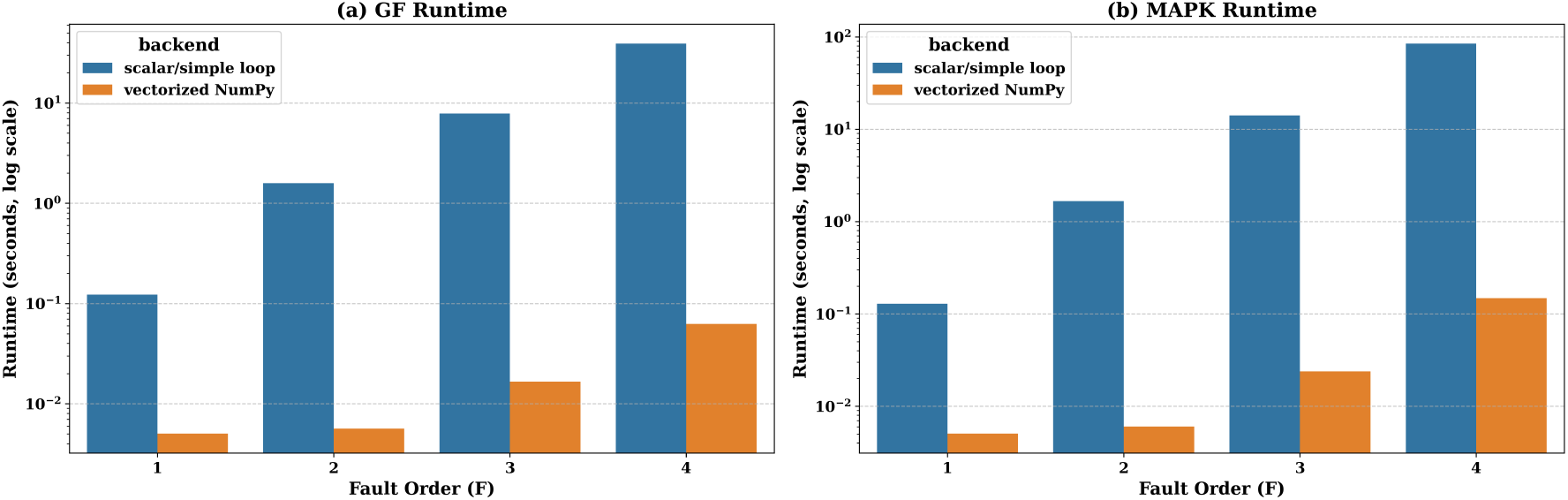
Runtime comparison between scalar/simple-loop execution and batched vectorized NumPy execution for GF and MAPK across fault orders

Table 17 summarizes the runtime benchmark results for scalar/simple-loop execution and batched vectorized NumPy execution. For the GF pathway, runtime increased from 0.1225s for 23 one-fault simulation jobs to 39.2112s for 8855 four-fault simulation jobs under scalar execution. The corresponding vectorized runtimes were 0.0050s, 0.0057s, 0.0166s, and 0.0625s for fault orders one through four, yielding speed-ups of 24.30×, 281.07×, 473.36×, and 627.18×, respectively. As shown in Fig. 16, the gap between scalar and vectorized execution widened substantially as the fault order increased.

**Table 17:**
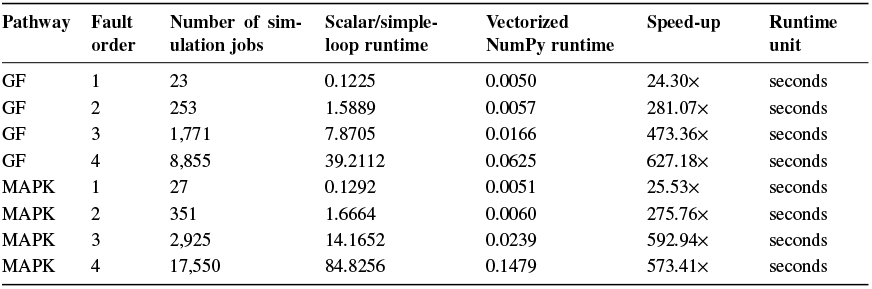
Runtime benchmark summary.

| Pathway | Fault order | Number of simulation jobs | Scalar/simple-loop runtime | Vectorized NumPy runtime | Speed-up | Runtime unit |
| --- | --- | --- | --- | --- | --- | --- |
| GF | 1 | 23 | 0.1225 | 0.0050 | 24.30× | seconds |
| GF | 2 | 253 | 1.5889 | 0.0057 | 281.07× | seconds |
| GF | 3 | 1,771 | 7.8705 | 0.0166 | 473.36× | seconds |
| GF | 4 | 8,855 | 39.2112 | 0.0625 | 627.18× | seconds |
| MAPK | 1 | 27 | 0.1292 | 0.0051 | 25.53× | seconds |
| MAPK | 2 | 351 | 1.6664 | 0.0060 | 275.76× | seconds |
| MAPK | 3 | 2,925 | 14.1652 | 0.0239 | 592.94× | seconds |
| MAPK | 4 | 17,550 | 84.8256 | 0.1479 | 573.41× | seconds |

A similar trend was observed for the MAPK pathway. Scalar execution increased from 0.1292s for 27 one-fault simulation jobs to 84.8256s for 17,550 four-fault simulation jobs. Vectorized execution reduced these runtimes to 0.0051s, 0.0060s, 0.0239s, and 0.1479s for fault orders one through four, corresponding to speed-ups of 25.53×, 275.76×, 592.94×, and 573.41×, respectively. Fig. 16 likewise shows that vectorized execution maintained low runtimes even as the number of simulation jobs increased dramatically.

These results show that the computational cost grows rapidly with fault order due to the increasing number of fault combinations to be evaluated. However, the vectorized NumPy implementation substantially reduced runtime across all scenarios, making largescale multi-fault probabilistic DBN analysis computationally practical for both pathways. The runtime comparison in Fig. 16 visually highlights the scalability advantage of the vectorized approach, particularly for the computationally demanding three and four-fault analyses.

## 6 Discussion

This study presents a probabilistic temporal DBN framework for evaluating multi-fault dysregulation and intervention response in the GF and MAPK signaling pathways. The results show that the framework can represent graded pathway burden as fault complexity increases, compare known-drug intervention vectors across fault orders, identify low-burden, high-efficiency candidates through Pareto ranking, and generate computational target hypotheses through a custom dual-target search.

### 6.1 Probabilistic DBN Interpretation of Multi-fault Burden

The selection of 00001 as the lowest-burden input vector for both GF and MAPK pathways provided a common baseline for all subsequent simulations. This input vector minimized the final output burden in both pathways, although the MAPK pathway retained a higher fault-free burden than GF. This difference is expected because the probabilistic DBN does not force the pathway into an exact all-zero output state. Instead, basal leakage, soft-logic propagation, and temporal persistence allow nonzero activation probabilities even in fault-free conditions.

The baseline multi-fault results in Fig. 3 and Table 12 show that encoded burden increased as the number of simultaneous faults increased. This confirms that concurrent pathway dysregulation produces a stronger downstream output burden than isolated single faults. However, the boxplots also show that fault order alone does not fully determine pathway severity. Some higher-order fault combinations produced lower encoded burden than expected, indicating that the location and interaction of faulty nodes are also important. This supports the need for combinatorial fault analysis rather than evaluating only representative or isolated fault cases.

Across all fault orders, MAPK showed a higher baseline encoded burden than GF. This suggests that, within the implemented DBN structure, MAPK is more sensitive to multi-fault perturbation. The larger output structure and additional downstream nodes in the MAPK model may contribute to this higher burden accumulation.

### 6.2 Known-drug Efficiency and Burden-aware Ranking

The known-drug efficiency analyses in Figs. 6-11 show that drug-vector performance followed structured patterns across both pathways and all fault orders. The single-fault heatmaps in Figs. 4 and 5 further demonstrate that intervention response depends on fault context: a drug vector may reduce burden effectively for some fault locations but less effectively for others. This supports the use of aggregate efficiency scores across all fault combinations of a given order.

The most consistent low-burden known-drug vector was U0126+LY294002+Temsirolimus. As shown in Table 13, this three-drug vector remained close to the maximum-efficiency six-drug vector across all pathways and fault orders. The maximum-efficiency vector, Lapatinib+AG1024+U0126+LY294002+Temsirolimus+Metformin, achieved the highest efficiency scores, but its improvement over the three-drug vector was small. Therefore, the results suggest that most of the achievable burden reduction can be obtained through downstream inhibition of MEK1, PIK3CA, and mTOR-associated signaling, without requiring the larger six-drug combination.

The Pareto analysis in Fig. 12 and Table 15 strengthens this interpretation. Efficiency increased sharply up to an intervention burden of three drugs, after which additional drugs produced smaller gains. Thus, the three-drug vector represents a favourable trade-off between intervention efficiency and intervention burden. This distinction is important because ranking candidates only by maximum efficiency would always favour larger combinations, whereas Pareto-based ranking highlights candidates that provide strong burden reduction with lower intervention complexity.

Although the absolute mathematical maximum efficiency was achieved by the six-drug vector Lapatinib+AG1024+U0126+LY294002+Temsirolimus+Metformin, the Pareto analysis indicates clear diminishing returns beyond an intervention burden of three. As shown in Fig. 12 and Table 15, the Pareto front exhibits a distinct knee around the three-drug combination U0126+LY294002+Temsirolimus. Beyond this point, adding further drugs produces only modest additional efficiency gains despite increasing the intervention burden. Therefore, within the proposed DBN framework, U0126+LY294002+Temsirolimus represents a favourable burden-aware trade-off: it achieves most of the encoded-burden reduction obtained by the maximum-efficiency vector while requiring only half the number of drugs. This makes it the most interpretable low-burden candidate identified by the known-drug analysis, although further biological and pharmacological validation would be required before any clinical interpretation.

The efficiency scores also decreased as fault order increased, as shown in Table 13 and the pathway-specific profiles in Figs. 10 and 11. This indicates that intervention becomes less effective as more pathway components are simultaneously dysregulated. Nevertheless, the same leading candidates remained stable across fault orders, suggesting that downstream pathway-targeting combinations retain relative importance even under more complex multi-fault conditions.

### 6.3 Custom Dual-target Candidates and Cross-pathway Differences

The custom dual-target analysis extended the intervention search beyond the predefined known-drug vectors and identified pathway-node pairs whose simultaneous inhibition produced high encoded-burden reduction. This analysis is important because it does not assume that the most effective intervention points must be limited to the targets of the seven known drugs. Instead, it searches the pathway topology directly and identifies node pairs that may behave as important control points in the probabilistic DBN model. The biological interpretation of the custom candidates becomes clearer when the DRG labels are mapped to the corresponding pathway nodes. As shown in Table 18, many of the top-ranked custom candidates involve nodes from the RAF-MEK-ERK axis, the IGF/IRS1-associated upstream branch, and the mTOR/S6K-associated downstream convergence region. This is biologically meaningful because these regions connect upstream receptor signals to transcription-factor and residual-protein outputs. Therefore, inhibiting these nodes can reduce signal propagation across multiple downstream routes.

**Table 18:**
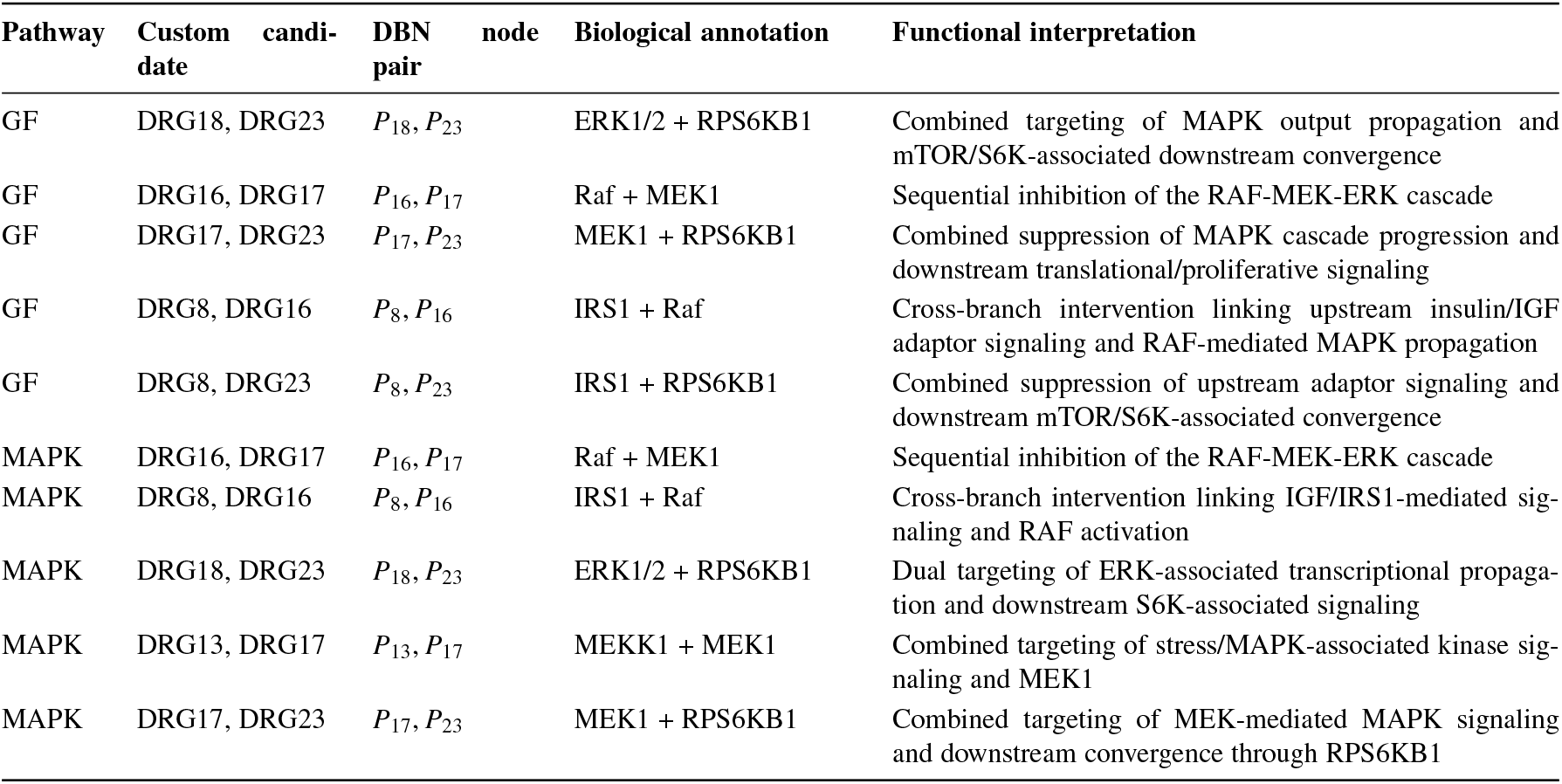
Biological annotation of top custom dual-target candidates.

In the GF pathway, the strongest custom candidate was [DRG18,DRG23], corresponding to P18/P23 or ERK1/2 and RPS6KB1. This pair is relevant because ERK1/2 is a central downstream component of the RAF-MEK-ERK cascade, while RPS6KB1 represents a downstream convergence point associated with mTOR/S6K-mediated signaling. Simultaneous inhibition of ERK1/2 and RPS6KB1 may therefore suppress both MAPK-mediated transcriptional propagation and mTOR/S6K-associated downstream output burden. This explains why [DRG18,DRG23] achieved the highest GF custom-target efficiency score in Fig. 13. Other high-performing GF custom pairs also support this interpretation. [DRG16,DRG17] corresponds to Raf and MEK1, representing two sequential components of the RAF-MEK-ERK cascade. [DRG17,DRG23] corresponds to MEK1 and RPS6KB1, combining inhibition of MAPK cascade progression with downstream S6K-associated signaling. [DRG8,DRG23] corresponds to IRS1 and RPS6KB1, linking an upstream IGF/IRS1-associated adaptor node with a downstream convergence node. These combinations suggest that high custom-target efficiency is achieved when the intervention simultaneously blocks signal entry or propagation and downstream output convergence.

In the MAPK pathway, the strongest custom candidate was [DRG16,DRG17], corresponding to Raf and MEK1. This result is biologically plausible because Raf and MEK1 are sequential nodes in the RAF-MEK-ERK cascade, one of the central signaling routes by which growth-factor stimulation is transmitted toward transcriptional outputs. Under multi-fault conditions, inhibition of only one node may not fully suppress signal propagation if alternative upstream activations remain active. Simultaneous inhibition of two consecutive nodes can create a stronger blockade within the cascade, explaining the high efficiency of [DRG16,DRG17] in Fig. 14.

The cross-pathway comparison in Fig. 15 further shows that some custom candidates performed well in both GF and MAPK, while others were more pathway-specific. [DRG16,DRG17] was strong in MAPK and also competitive in GF, suggesting that the RAF-MEK axis is a robust intervention region across both pathway models. In contrast, [DRG18,DRG23] produced the highest GF score but a lower MAPK score, indicating that the effectiveness of ERK1/2 and RPS6KB1 co-inhibition depends on the pathway-specific topology and output structure.

Overall, the custom dual-target results suggest that high-performing intervention pairs tend to involve either sequential blockade within a major signaling cascade or combined suppression of upstream/adaptor signaling and downstream convergence. However, these candidates should not be interpreted as validated drug combinations. They are computational target hypotheses generated by the DBN model. Their relevance lies in identifying pathway regions that may be worth further investigation through druggability assessment, biological perturbation experiments, and pharmacological validation.

### 6.4 Computational Scalability

The runtime benchmark in Fig. 16 and Table 17 shows that batched vectorized NumPy execution substantially reduced computational time compared with scalar/simple-loop execution. This improvement became especially important in higher-order fault scenarios, where the number of fault combinations increased rapidly. For four-fault simulations, the vectorized implementation produced speed-ups greater than 500× in both pathways.

This scalability is important because the framework requires repeated simulations across fault combinations, 128 known-drug vectors, and custom dual-target candidates. Without vectorized execution, the higher-order multi-fault analyses would become computationally expensive. The runtime results, therefore, show that batched simulation is a practical requirement for extending probabilistic DBN analysis to larger pathways, broader intervention libraries, or additional parameter settings.

### 6.5 Limitations and Future Work

As with any computational modeling study, several considerations should be kept in mind when interpreting the results, while also recognizing the strengths and contributions of the proposed framework. First, the model is entirely in silico. The efficiency scores quantify reductions in encoded DBN output burden rather than experimentally measured drug response. Consequently, the known-drug vectors and custom target pairs identified in this study should be viewed as computationally prioritized candidates that provide a focused basis for subsequent experimental investigation. Second, the drug-action representation is intentionally abstracted to enable systematic exploration of large intervention spaces. Known drugs are modeled through mapped pathway targets and probabilistic intervention rules. While this approach effectively supports comparative analysis across pathways and fault conditions, it does not currently incorporate factors such as dose response, pharmacokinetics, pharmacodynamics, toxicity, off-target effects, tissue specificity, or patient-specific variation. Likewise, intervention burden serves as a computational measure of intervention complexity rather than a direct clinical toxicity metric.

Third, the custom dual-target candidates are not intended to represent existing drugs. Instead, they identify pathway-node pairs whose simultaneous inhibition produces favorable outcomes within the DBN framework. These findings provide potentially valuable hypotheses for future drug-discovery and target-validation efforts, where biological feasibility, druggability, and experimental effectiveness can be further assessed.

Fourth, the results are naturally influenced by the selected pathway topology, rule definitions, node mappings, and DBN parameter settings. Although the consistency of several leading candidates across pathways and fault orders supports the robustness of the framework, additional parameter-sensitivity analyses could further strengthen confidence in candidate rankings under alternative assumptions. Future extensions may also incorporate larger signaling networks, feedback mechanisms, data-driven parameter estimation, drug synergy models, and experimental validation.

From a broader perspective, future research should focus on both computational expansion and biological validation. The low-burden known-drug vector U0126+LY294002+Temsirolimus is particularly noteworthy because it consistently emerged as a strong burden-3 candidate across pathways and fault orders. Evaluating this combination experimentally, alongside the identified custom target pairs, would provide valuable insight into the predictive utility of the proposed framework. Such developments would further enhance the applicability of the methodology while preserving its key strength: the ability to perform systematic multi-fault and multi-intervention analysis within a unified computational framework.

## 7 Conclusion

This study presented a probabilistic temporal Dynamic Bayesian Network-based framework for analyzing multi-fault dysregulation and intervention response in the GF and MAPK signaling pathways. By propagating activation probabilities over discrete time steps, the framework enabled graded assessment of pathway behaviour using encoded burden and intervention efficiency scores across one, two, three, and four-fault scenarios.

The results showed that encoded pathway burden increased with the number of simultaneous faults in both pathways, with MAPK consistently showing higher burden than GF. Known-drug analysis identified U0126+LY294002+Temsirolimus as the most consistent low-burden intervention vector across both pathways and all fault orders. Although the six-drug vector Lapatinib+AG1024+U0126+LY294002+Temsirolimus+Metformin achieved the highest mathematical efficiency, Pareto analysis revealed diminishing returns beyond a burden of three drugs. Thus, the three-drug vector represented a favourable burden-aware trade-off, achieving most of the encoded-burden reduction while requiring only half the intervention burden.

The custom dual-target analysis further identified high-impact computational target hypotheses, including ERK1/2+RPS6KB1 in GF and Raf+MEK1 in MAPK. These candidates highlight pathway regions where simultaneous inhibition may strongly reduce downstream burden, although experimental validation and druggability assessment are required before any therapeutic interpretation. Runtime benchmarking also showed that batched vectorized NumPy execution substantially improved scalability, making higher-order multi-fault simulations computationally feasible.

Overall, the proposed probabilistic DBN framework provides a scalable and interpretable approach for pathway-level multi-fault analysis and intervention prioritization. It identifies robust low-burden known-drug candidates, suggests biologically meaningful custom target pairs, and provides a foundation for future extensions involving larger pathway maps, data-driven parameterization, broader drug libraries, and experimental validation.

